# Recon8D: A metabolic regulome network from oct-omics and machine learning

**DOI:** 10.1101/2024.08.17.608400

**Authors:** Ryan Schildcrout, Kirk Smith, Rupa Bhowmick, Yuntao Lu, Suraj Menon, Minali Kapadia, Emily Kurtz, Anya Coffeen-Vandeven, Srikar Nelakuditi, Sriram Chandrasekaran

## Abstract

To explore multiomic regulation of the metabolome, we used machine learning to predict metabolomic variation across ∼1000 different cancer cell lines with matched omics data from 8 biomolecular classes: genomics (copy-number and mutations), epigenomics (histone post-translational modifications (PTMs) and DNA-methylation), transcriptomics and RNA splice variants, non-coding transcriptomics (miRNA and lncRNA), proteomics, and phosphoproteomics. Overall, the metabolome is tightly associated with the transcriptome, with coding and non-coding RNAs emerging as top predictors. Peripheral metabolites are predictable via levels of corresponding enzymes, while those in central metabolism require combinatorial predictors in signaling and redox pathways, and may not reflect corresponding pathway expression. We reconstruct multiomic interaction subnetworks for highly predictable metabolites, and YAP1 signaling emerged as a top global predictor across 4 omic layers. We prioritize predictive multiomic features for single-cell and spatial metabolomics assays. Top predictors were enriched for synthetic-lethal interactions and synergistic combination therapies that target compensatory metabolic modulators.

## Introduction

Variation in metabolite levels between cells may arise due to differences in regulation at the transcriptional, post-transcriptional, translational, and post-translational levels^1,2^. Understanding metabolic regulation in a holistic manner remains challenging due to the complex interconnected nature of metabolic networks, comprising thousands of reactions, that are controlled by an equally complex web of regulatory interactions^3^. In addition to thermodynamics and nutrient availability, intracellular metabolite levels are driven by enzyme activity, which is the basis for the metabolome’s relationship with proteogenomic networks. Because of their wide availability, transcriptomics or proteomics data have been used to predict the metabolome with limited success^4–7^. However, these studies focus on a limited set of omics data and do not examine the relative influence and predictive power of each omics class on the metabolome^8–18^. Further, it is unclear if certain metabolites are more predictable than others.

Here, we leverage natural metabolic variation across hundreds of cancer cell lines to infer a multiomic predictor network of the metabolome (Recon8D) using machine learning (ML). Our approach utilizes 10 omics inputs from 8 biomolecular classes from the Cancer Cell Line Encyclopedia (CCLE), allowing for direct comparison of each class on predicting metabolite variation. We hypothesized that the extent of predictability is indicative of the strength of relationship with the metabolome. Similar assumptions have been used in numerous studies to assess relationships between transcription factors and target genes^19–22^. Although many metabolic regulatory relationships have been documented, they represent a small fraction of the number of interactions that likely exist in biological systems. ML algorithms can fill this gap by learning relationships from experimental data. Our approach uncovered numerous mechanistic relationships and identified significant multiomic predictors of metabolomic variation that could serve as markers for either individual metabolites or the overall metabolome for *in vivo*, single-cell or spatial omics studies that require limited measurements^23–26^. Finally, our discovery of synthetic lethal interactions among top metabolite predictors can help design combination therapies by targeting compensatory mechanisms.

## Results

### Recon8D model construction using machine learning

Analogous to gene regulatory inference with ML, wherein the expression value of each gene is taken as the learning objective, here we use metabolite level variation as the prediction task in a regression problem using multiomic data as input^21,27^. Each omics class is used individually to predict the metabolome, enabling rational comparisons of predictive capabilities across omics. The output is an ‘influence’ network model that implicitly captures regulatory events at the transcriptomic, proteomic or post-translational levels. Our approach used omics from the CCLE, which include biomolecular assays from up to ∼1,000 cancer cell types^28^, to predict the corresponding metabolome. We used 8 biomolecular classes: genomics – 1. copy number variation (CNV) and 2. mutations, epigenomics – 3. DNA methylation and 4. histone PTMs, transcriptomics – 5a. coding transcripts and 5b. RNA splicing, non-coding transcriptomics – 6a. miRNA and 6b. lncRNA, 7. proteomics, and 8. phosphoproteomics to train ML models for predicting relative levels of each metabolite across matched cell lines (Figure 1, S. Table 1).

**Figure 1:**
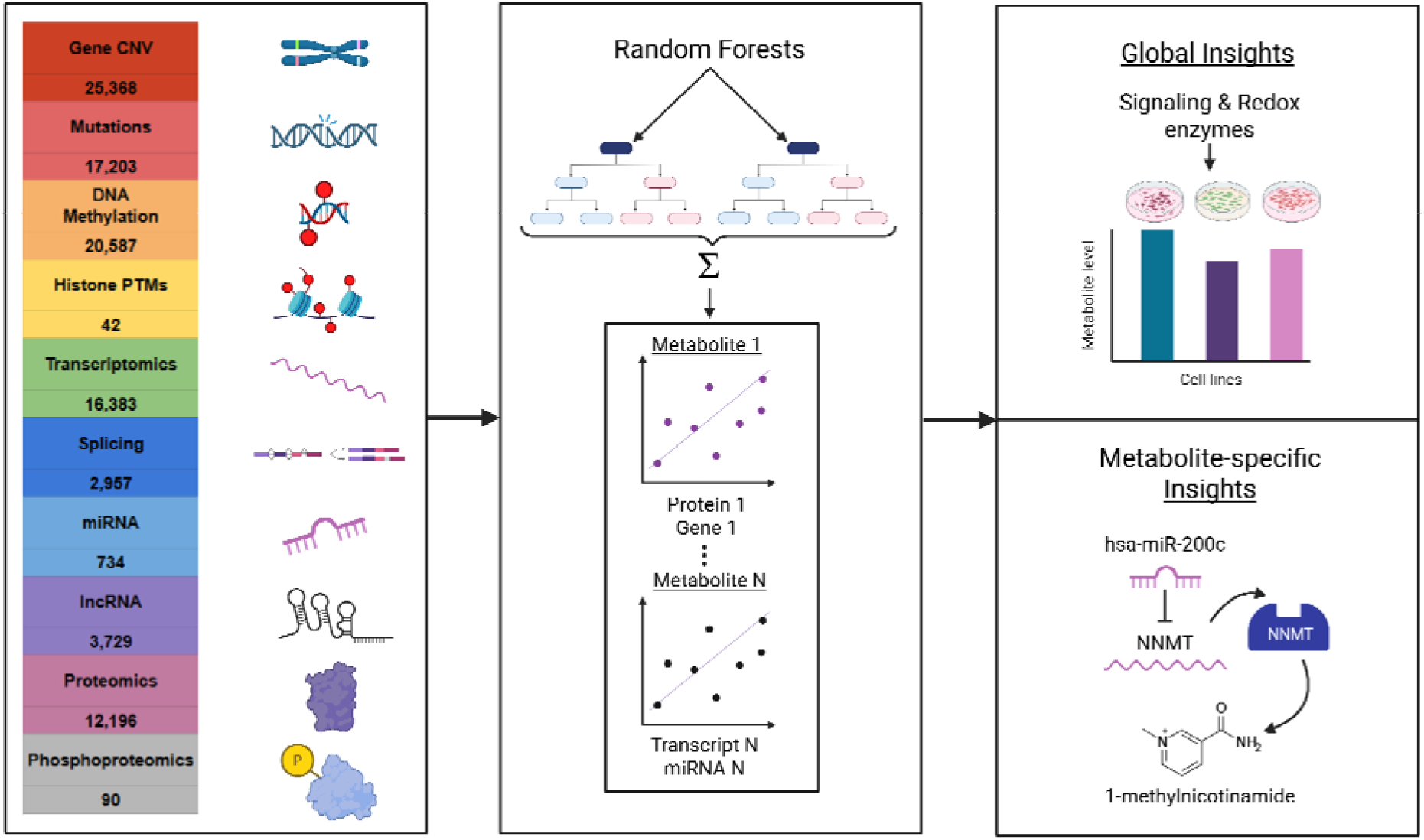
Graphical overview of our approach to predict the metabolome from multiomics and inferring top global and metabolite-specific predictors. Each omics class is utilized in isolation to train a predictive model for each metabolite in a metabolomic panel, and the accuracies and feature importances thereof are used to reconstruct Recon8D metabolic regulatory networks.

Random forest regression ML algorithms were used to generate metabolomic predictions, as they generally outperformed other ML methods based on cross-validation in our benchmarking (S. Figure 1). Random forests have also outperformed other methods in the inference of gene regulatory networks in prior studies^21,27^. We further make use of inherent characteristics of random forests to obtain the importance of each feature in prediction tasks. Higher feature importances indicate a greater possibility that a gene or protein regulates the target metabolite. Separate random forest models were trained iteratively for each of 225 metabolites in the CCLE metabolome using each of the omics features in an input set to create a total of 2,025 models (Figure 2A, 2B). Model performance was evaluated using cross-validation, and significantly predictable metabolites were identified based on a Bonferroni-corrected P value of 0.05 to account for false discovery rate (Figure 2C). We aimed to avoid bias in our models by employing a wide array of omics without incorporating information from previously known molecular interactors, as some omics classes are studied far more in depth than others. This framework enables rational comparisons across omics classes regardless of previous knowledge on their regulatory influence.

**Figure 2.**
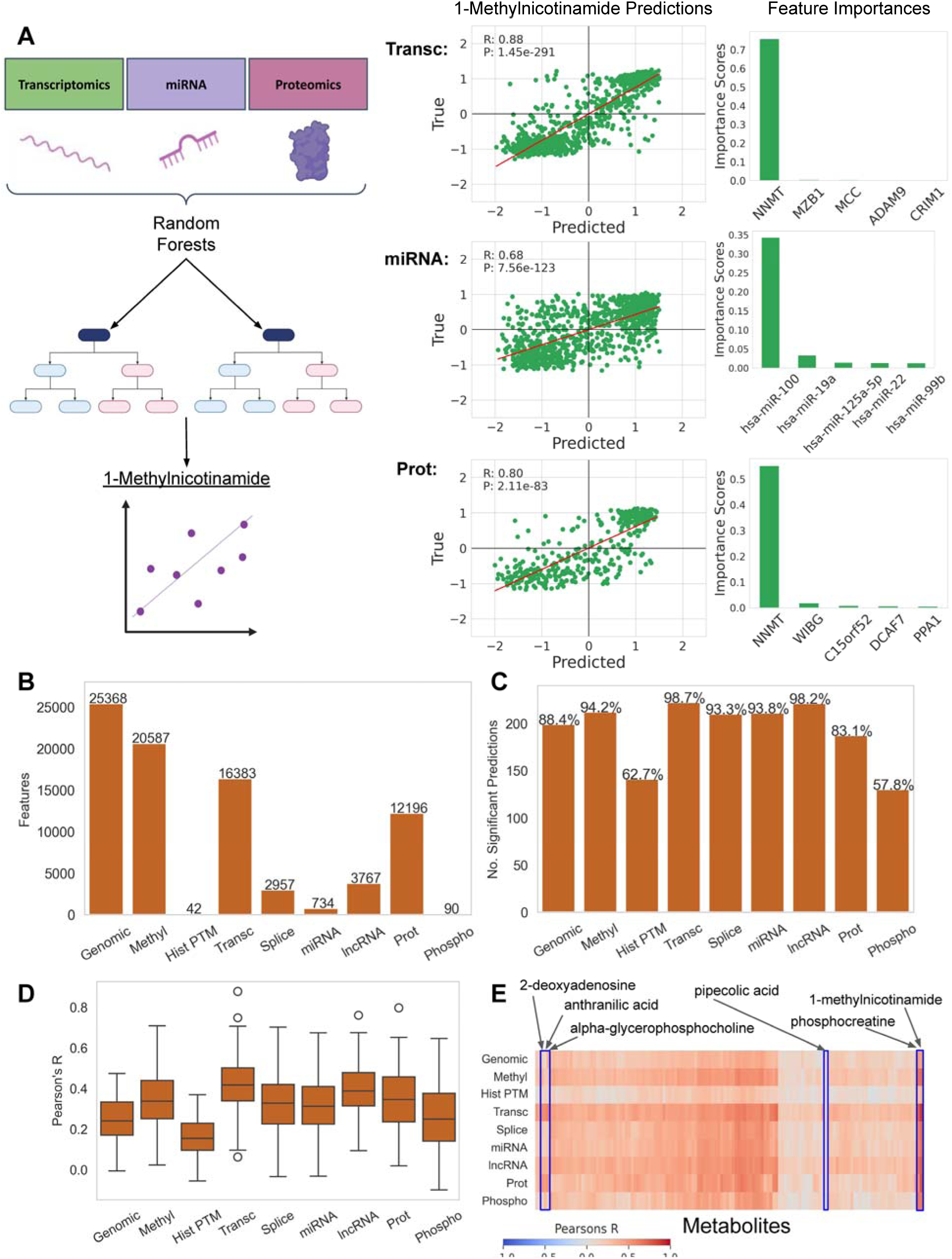
Random forest model performance indicates transcriptomics, DNA methylation, and non-coding transcriptomics are accurate predictors of metabolomic variation. **A.** Representative examples of 1-methylnicotinamide predictions using transcriptomic, miRNA, and proteomic inputs. Each omics class is used individually to predict relative 1-methylnicotinamide levels, which yields a predicted metabolite variation array which can be compared to the true values. Scatter plots of predicted vs true values in cross-validation help visualize the accuracy of each model. Finally, feature importance analysis identifies the most influential features on the model output. In this case, the model identified NNMT, the enzyme involved in 1-methylnicotinamide synthesis, as the top feature predictive of 1-methylnicotinamide. **B.** Number of features in each input dataset. **C.** Number of significantly predicted (FDR p-value < 0.05) metabolites per input omics class for each of 225 metabolites measured in the CCLE panel, also represented as a percentage out of 225. **D.** Box plots showing the distribution of Pearson’s R values for predicted vs true values from each omics model type over 225 metabolite models. **E.** Clustered heatmap showing correlation values for all non-lipid metabolites (columns) with select metabolites labeled.

### ML model performance indicates metabolic variation reflects transcriptional differences between cell lines

Across all ML algorithms and validation procedures, transcriptomics consistently generated the most accurate predictions (Figure 2, S. Figures 1, 3, and 4). While histone PTMs and phosphoproteomics were not as accurate as other omics classes, they had fewer features by an order of magnitude, showcasing that small feature sets can still impart meaningful predictions. The ability of each omics class to make accurate predictions independent of feature set size instills confidence in the hypothesis that the ML models are picking up mechanistic relationships and are not overfitting to the data. For example, with the seventh-most number of features out of 9 omics classes, miRNA inputs produced the fourth-most significant predictions. In contrast, genomics (CNV) had the most features, yet only the sixth-highest predictive performance. We further validated this assumption with benchmarking models trained on the top 40 features per dataset identified using holdout validation with a training set to testing set ratio of 75 to 25 (S. Figure 1). The transcriptomic and miRNA data achieved high accuracy (96% of metabolites predicted with FDR p-value < 0.05) while only using 40 features. The top 40 features can thus potentially serve as a reduced set of molecular markers that can be measured for reconstructing the metabolome (S. Table 2).

Box plots and clustered heatmaps of Pearson’s correlations (model predictions versus true relative metabolite levels) illustrate the differences in predictive power across omics inputs (Figure 2D, E). While box plots help visualize predictability across the metabolome, the heatmap of individual models is useful for identifying metabolites that are differentially predictable with certain omics inputs. While some metabolites are highly predictable across omics layers, such as 1-methylnicotinamide and phosphocreatine, others have stark differences in predictability. For example, anthranilic acid, 2-deoxyadenosine, and alpha-glycerophosphocholine predictions were more accurate with transcriptomic and proteomic inputs. This suggests that individual metabolites can be influenced preferentially at particular omics levels. Conversely, some metabolites, such as pipecolic acid, do not have high correlations with any model, suggesting that their mode of regulation might not be sufficiently captured in these data.

The ability to quantify associations with the metabolome across omics classes is vital for prioritizing the study of novel regulators. Globally, model results indicate that non-coding RNAs are highly associated with metabolomic variation across cell lines as compared to other omics classes, highlighting their importance for further study in context with metabolic regulatory networks.

### Control and validation experiments show consistency and robustness in predictive capability

To account for confounding factors that may drive spurious correlations, such as tissue type, culture conditions, and tumor sample origin, several control experiments were conducted. We trained models using omics subsets separated by growth type (adherent and suspension), cancer lineage (lung and hematopoietic/lymphoid tissue, as well as all lineages except lung or hematopoietic/lymphoid tissue), and by tumor sample origin (primary and metastatic) (S. Figure 3). Out of the 8 control experiments conducted, confidence scores were calculated for each feature (0-8) depending on controls where the feature appeared in the top 20 (Methods). The goal of this system is to assess the ability of the models to identify regulator-metabolite relationships rather than assess overall accuracy. Top features discussed throughout are shown with corresponding confidence scores (CS: 0-8), and those discussed in the text have a CS of at least 2, implying that they were observed in the overall model and at least two other control conditions. While the raw accuracies of the models in these experiments are lower due to loss of observations (cell lines) per dataset, thereby limiting training data, similar trends in the predictive power of each omic subset are observed across controls. Models controlled for tumor origin show a high level of concordance in accuracy and feature importance. Overall, transcriptomics had the highest or second-highest predictive accuracy across all omics models in all control experiments.

To assess the robustness of the top features in independent datasets, validation experiments were conducted where top features from CCLE-trained models were compared to corresponding relationships between features and metabolites in the NCI60 cancer cell line panel^29^, similar to the control experiments described above. Random forest models were trained on NCI60 genomics (CNV), transcriptomics, miRNA, proteomics, and phosphoproteomics data, and top features from the CCLE models were compared to the top features from the NCI60 models (Methods). CCLE-identified features showed significantly higher feature importance scores in the NCI60 data as compared to the whole feature set (S. Tables 3-4). To further assess the generalizability of this framework to single-cell datasets, wherein the metabolome is challenging to measure, we performed a similar analysis described above with single-cell transcriptomics of the cell cycle and compared with matched metabolome from synchronized cells^30,31^. CCLE-identified features show significantly higher feature importance scores in the single-cell transcriptomics data than expected out of random chance (S. Table 5). These experiments demonstrate generalizability of the feature-metabolite associations from CCLE-trained models across cell line panels.

### ML Feature Importance Analysis reveals global and metabolite-specific predictors

For each metabolite, we looked at top predictors from each omics-based model based on feature importance scores (Methods). The transcriptomics model not only had high predictive power, but also recalled several known mechanistic metabolite-gene relationships. For some metabolites, the mRNA transcript of the corresponding enzyme that synthesizes the metabolite was the top predictor. These include choline O-acetyltransferase gene (CS: 4) for the metabolite acetylcholine, choline dehydrogenase (CS: 4) for betaine, cystathionine beta-synthase (CS: 7) for cystathionine, and glycerol 3-phosphate dehydrogenase (CS: 7) for alpha-glycerophosphate. For other metabolites, the consumption or catabolic degradation was more predictive of their levels.

For example, cytidine deaminase (CS: 6) for cytidine, and monoamine oxidase A (CS: 2) for serotonin. Several metabolites’ corresponding transporters also featured as top predictors, including SLC6A6 (CS: 8) for taurine and SLC6A8 (CS: 8) for phosphocreatine. Nicotinamide and tryptophan metabolism shared several top predictors, suggesting shared regulation. Nicotinamide phosphoribosyltransferase (NAMPT) (CS: 4) was a top predictor of NAD. Indoleamine 2,3-dioxygenase (IDO) (CS: 6) and tryptophan 2,3-dioxygenase (CS: 5) were most predictive of kynurenine. Anthranilic acid was predicted by kynureninase (CS: 6), and both methylnicotinamide and niacinamide were predicted by nicotinamide N-methyltransferase (NNMT) (CS: 8 and 4, respectively).

The proteomics model also recapitulates several known relationships between metabolites and specific enzymes that impact their levels. For example, the top predictor in the proteomics model for cystathione was cystathionine beta-synthase (CBS) (CS: 5), and the top predictor for phosphoenol pyruvate was pyruvate kinase (PKM) (CS: 3). Thymine and uracil’s top predictor was dihydropyrimidine dehydrogenase (DPYD) (CS: 2 and 5, respectively) - the initial and rate-limiting enzyme in uracil and thymidine catabolism. Similarly, malic enzyme (ME2) (CS: 4), part of the malate-aspartate shuttle, was a top predictor of aspartate levels, while glutaminase (GLS) (CS: 3) was a top predictor of glutamine.

While many metabolites were predicted preferentially by a single top feature, others were predicted by multiple features, suggesting that they might be involved in multiple pathways and may be regulated combinatorially. Such metabolites will be harder to predict using transcriptomic variation of a single gene. For example, citrate is an intermediate in multiple core metabolic pathways, and its concentration remains low in cancer due to its rapid conversion into acetyl-CoA and oxaloacetate^32^. As opposed to metabolites such as taurine where there is one clear regulator, citrate’s feature importances show a more balanced regulatory influence among several top features (Figure 3D). The top predictor for taurine was SLC6A6 (CS: 8), the taurine transporter, and was shown by our models to have an extremely high influence in predicting relative taurine levels as compared to even the second most important feature by Shapley analysis (Figure 3D). However, citrate’s feature importance plot does not show one clear top predictor, but rather an array of features that appear to impact relative citrate levels similarly. In this way, ML can uncover complex metabolic control by multiple inputs in addition to strong associations between single features and metabolites. Interestingly, metabolites that had a distinct single predictor like anthranilic acid, N-carbamoyl-beta-alanine, and adenosine were also more generalizable to NCI60 (S. Table 3). This may also be related to the limited ability of ML to tease out complex combinatorial relationships using small data such as the NCI60 panel.

**Figure 3.**
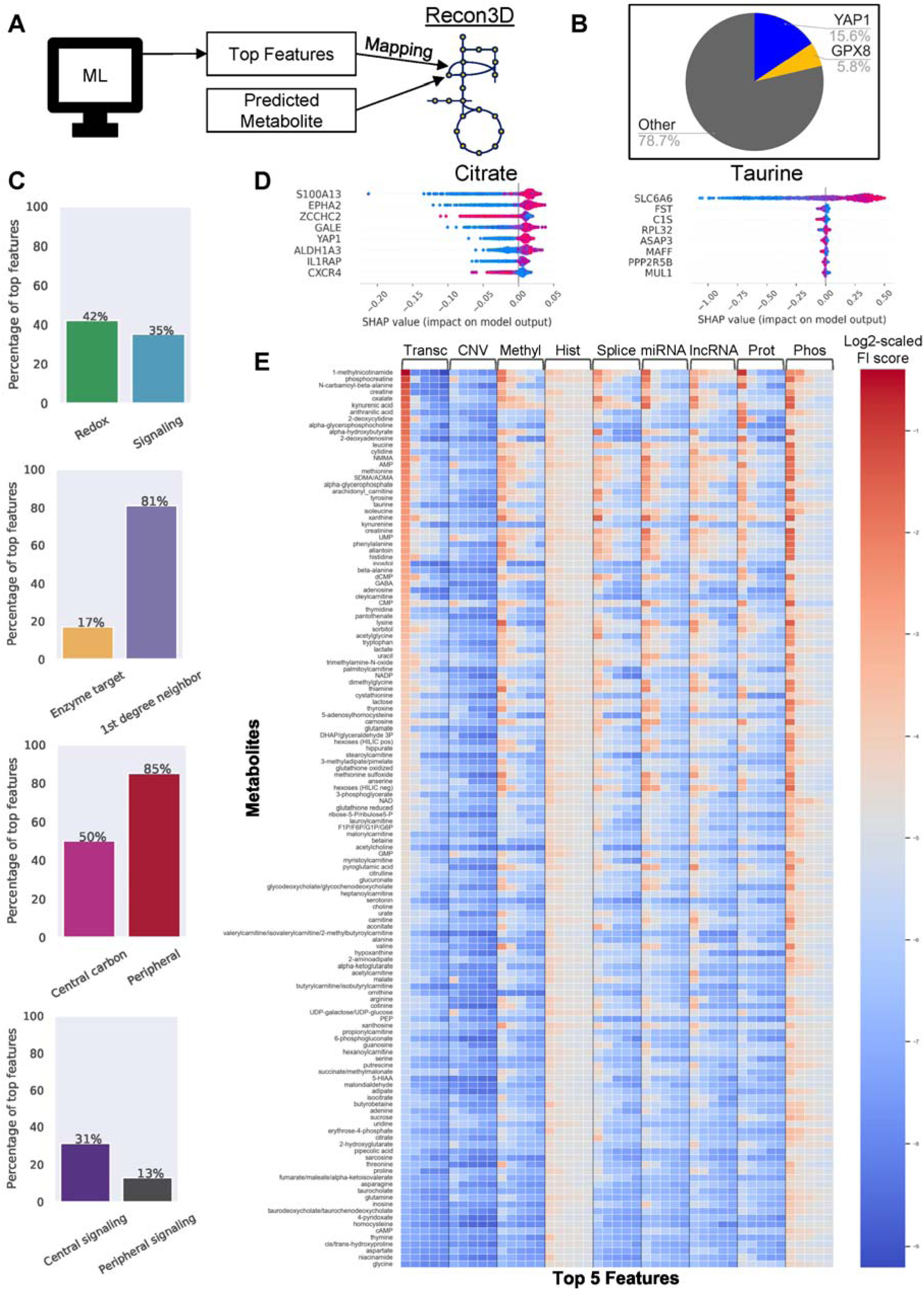
Global insights across all 225 ML models reveal distinct patterns of predictability. **A.** Workflow for topologically mapping features to Recon3D. Top 1, 2, and 5 features are identified from ML, and are assessed for overlap with genes present in the reactions for its predicted metabolite (both the enzyme target and the 1st degree neighbor). **B.** Pie chart showing the percentage (of 225) of top features from transcriptomic inputs. YAP1 and GPX8 account for 15.6% and 5.8% of top 1 features, respectively. **C.** Percentage of top 1 features from biologically relevant feature subsets using transcriptomics inputs. 42% (94) of top features (out of 225 metabolite models, metabolic gene-only) are redox enzymes, and 35% (79) of top features (out of 225 metabolite models, whole-data) are signaling enzymes. 17% (16) of top features (out of 95 metabolites that overlapped between CCLE and Recon3D) were involved in the primary reaction for their respective metabolite, and 81% (77) of top features (out of 95 metabolites) were involved within the 1st neighboring reaction for their respective metabolite. 50% (5/10) top features from the central carbon metabolites and 85% (72/85) of peripheral metabolites (from 95 total metabolites that overlapped between CCLE and Recon3D) were involved within the 1st neighboring reaction for their respective metabolites. Lastly, out of 136 non-lipid metabolite models, 31% (5/16) of top features were signaling enzymes for central carbon metabolites, and 13% (15/120) of top features were signaling enzymes for peripheral metabolites. **D.** Contrasting Shapley importance features for citrate and taurine from transcriptomics models. While the taurine transporter SLC6A6 is clearly shown to be the most important feature in predicting relative levels of taurine, citrate does not have one clear top predictor. In this way, feature importance analysis of citrate shows high order, complex influence by multiple features. **E.** Heatmap of log2-scaled feature importance scores for 136 non-lipid metabolites. The top 5 features for each omics class are shown for every metabolite in order of transcriptomics feature importance (descending). This shows a set of metabolites that have 1 strong predictor (top of heatmap) and those metabolites, primarily related to central metabolism, that have multiple predictors (bottom of the heatmap).

Analysis of genomic mutations data revealed several well-known associations between genetic regulators and metabolite variation in cancers. Due to its binary structure, mutation data was used to predict metabolite variation using support vector machines (SVMs) (S. Figure 1). While relatively less accurate than other omics across the metabolome, the mutation ML models nevertheless recapitulated known associations between important cancer markers and the metabolome. Out of the 107 metabolites predicted significantly with mutation data, mutual information calculations between mutation features and metabolites revealed a handful of critical cancer-related genes consistently appearing in the top 20 features across all metabolites (S. Table 6). For example, TP53 appeared in the top 20 features 20 times, APC 19 times, CDKN2A 17 times, PTEN 16 times, RB1 14 times, and MUC16 12 times. TP53 was a top feature for several nucleotides and APC was associated several amino acids and lipids, consistent with their known effects on cancer metabolism (S. Table 7).

The ML models also have the ability to identify potential novel associations between features and metabolites. While the top feature for 7 out of 10 of the most accurately predicted metabolites from transcriptomics was found to be mechanistically connected to its respective metabolite, the remaining 3 metabolites–oxalate, kynurenic acid, and alpha-hydroxybutyrate–had YAP1 (CS: 3, 3, and 3, respectively), a signaling protein, as the top predictor. Oxalate has been found to be associated with YAP1^33^, but the association between kynurenic acid or alpha-hydroxybutyrate and YAP1 has yet to be established.

Comparison of predictive capabilities between omics is particularly useful in the case of lncRNAs, where very few features have known functions, yet show tight associations with the metabolome. Overall, lncRNAs produced highly accurate metabolome predictions, generally second-best behind transcriptomics of protein coding genes. Interesting patterns emerged in the lncRNA data, such as MIR3936HG being the top feature for carnitine, acetyl-, propionyl-, malonyl-, butyryl-, and valeryl-carnitine (CS: 7, 7, 6, 6, 4, and 8, respectively). Top lncRNA features with high confidence scores (>5) prioritized for further study in context with their predicted metabolites are provided in the supplementary materials (S. Table 8).

The ML models also revealed long range metabolite-enzyme interactions. Top features in the proteomics model for glutathione, a key redox metabolite, included SLC7A11 (CS: 3) and ALDH3A1 (CS: 3). SLC7A11 is a cysteine and glutamate transporter (xCT), and its activity has been shown to influence glutathione levels^34^. Aldehyde dehydrogenase protects cells from oxidative stress and consumes redox cofactors. Interestingly, SLC7A11 and ALDH3A1 also form a synthetic lethal pair, suggesting that they may compensate for each other’s activity in maintaining redox balance^35^.

Interestingly, the top predictors for a majority of the metabolome across various omics data involved either signaling proteins like YAP1 or redox enzymes such as GPX8 (Figure 3B and C). YAP1 was among the top 2 predictors in the transcriptomics model for 45 metabolites including numerous amino acids and nucleotides, while GPX8 was among the top 2 predictors for 23 metabolites including various lipids. This suggests that the combination of growth signaling and redox homeostasis influences a significant part of the metabolome.

Global feature importances from control experiments also identified features integral to specific cellular states. For example, the ML models built using primary or metastatic cell lines revealed distinct miRNA predictors that are known drivers of tumor phenotypes in each condition. miR-142-5p, which has been shown to limit cell invasion and migration^36^, was the second most important global feature from the miRNA models and the most important in the primary tumor controls. miR-142 was also a top global feature in the DNA methylation data, 4th in whole-data models and 3rd in primary controls, but is not a top predictor in the metastatic ML models. In the metastatic samples, miR-142 was ranked 233rd out of 734 miRNAs, showing that its relation to metabolite level variation is blunted in the metastatic state. Additionally, miR-181a, which is known to promote proliferation and invasion^37^, is the 44th most important feature in the primary tumor control models versus 17th in the metastatic controls, highlighting its association with metabolism in metastatic cells in particular.

### Top predictors are close topologically to the corresponding metabolites

To further substantiate the ability of ML to identify metabolic regulatory interactions, we mapped features from genomic, DNA methylation, transcriptomic, and proteomic models to reactions present in the Recon3D model (Figure 3A, Methods). Recon3D is a metabolic network model built on biochemical reactions curated from literature^38^. It is a comprehensive collection of known metabolic reactions in human cells, making it a prime candidate for systematically validating ML models and uncovering topological relationships between top features and their respective predicted metabolites. To ensure overlap with Recon3D, ML models were re-trained on the Recon3D metabolic gene subset. The resulting metabolic subset ML models showed similar accuracy to whole-data models (S. Figure 4B and C). Feature importances from metabolic gene-only models and whole-data models were also conserved significantly (S. Figure 4D and E).

The top 5 features for each metabolite from all the metabolic ML models displayed a substantial number of overlaps (92-96%) with either the corresponding reactions directly associated with each metabolite or their first-degree neighbors (S. Table 9). This suggests an arms-length or 1-step influence strategy wherein the top predictors for ∼95% of the metabolites are within 1 step of the reactions involving those metabolites.

While the top features were on average topologically closer to the corresponding metabolite, consistent with prior studies^6,39^, we found distinctions for metabolites in peripheral or central metabolism (Figure 3C). For example, in the transcriptomics ML models, the top features for only 50% of the metabolites in central carbon metabolism overlapped with either the corresponding reaction or first-degree neighboring reactions. In contrast, the top features for 85% of the metabolites in peripheral metabolism were direct or first-degree neighbors, suggesting that peripheral metabolites are more influenced by direct enzymatic regulation, as opposed to those in central carbon metabolism, which are likely influenced more by signaling (Figure 3C).

To further explore if signaling is a larger player in central carbon metabolism than peripheral metabolism, we determined if top features in each category were signaling enzymes for all non-lipid metabolites. We found that 31% of metabolites in central carbon metabolism had signaling enzymes as the top feature compared to only 12.5% for peripheral metabolites (Figure 3C). These findings suggest that central carbon metabolism is influenced to a greater extent by signaling than peripheral metabolism.

### YAP1 and GPX8 are globally important features across the metabolome

After identifying the most influential features locally for individual metabolite models, we next pinpointed features that were globally important across the metabolome using the average rank of the features for all 225 metabolite models (Methods, S. Table 10). These results were then scaled by Z-score and top features for each input omics class were identified. Important features of interest were checked for overlap between whole-data models and controls for robustness.

Some top features were several standard deviations above the mean for certain omics classes, such as YAP1, GPX8, and SCD in the transcriptomics models, with Z-scores of 69, 61, and 59, respectively. Well known metabolic regulators such as mTOR also featured in the global top regulators. mTOR phosphorylation was a top 5 feature for several metabolites in central and nucleotide metabolism including oxalate, succinate, xanthine, and phosphogluconate; similarly, phosphorylation of Rictor, which interacts with mTOR was a top feature for amino acids including arginine, consistent with their role in amino acid sensing and regulation^40^.

Histone PTMs are increasingly being recognized for their role in sensing and regulating metabolism^41–43^. The CCLE epigenomics data measures total PTM levels and lacks gene-level resolution. Total levels of H3K9 trimethylation and H3K27 di-& trimethylation were the top features in histone PTMs; these are well known repressive marks, and importantly, are also sensitive to the metabolic state of the cell. Histone methylation is sensitive to the level of one-carbon metabolites like SAM and one-carbon sink metabolites such as methyl nicotinamide^42^.

Interestingly, top global predictors showed consistency across omics classes. The most intriguing example of this was YAP1, which was found to be the most important global feature for transcriptomics and phosphoproteomics, as well as the seventh for DNA methylation. YAP1 is a coactivator in Hippo signaling, which is essential for growth signaling, oncogenic adaptations, and the epithelial mesenchymal transition (EMT)^44^. YAP1 was the most important feature for 35 metabolite models using transcriptomics, 2 using proteomics, 5 using DNA methylation, and 132 using phosphoproteomics. Other examples of cross-omic consistencies for top features were GPX8 (transcriptomics and proteomics), miR142 (methylation and miRNA), EPHA2 (transcriptomics and RNA splicing), and ERBB2 (DNA methylation and proteomics). Not only do different omics display conserved global regulators, but globally important features have established relationships. For example, the top global feature from RNA splicing, EPHA2, has been shown to induce accumulation and activation of YAP1, subsequently promoting glutamine metabolism^45^.

To validate the association between YAP1 transcripts and the 35 metabolites for which it was the top predictor, we used the DepMap Data Explorer to quantify the relationships between two isoforms of YAP1 protein and each metabolite with linear regression (S. Table 11)^46^. We found that most associations were statistically significant (FDR p-value < 0.05), thus increasing confidence that YAP1 acts as an important predictor for a wide array of metabolites.

### Construction of multiomic metabolite regulome networks for top predictive metabolites

The Recon8D network, containing the top 10 predictors identified from random forest feature importance scores from each of the omic layers for all 225 metabolites is available in S. Table 12 and on the <u>GitHub page</u>. To understand how top omics features interact with one another as well as with the metabolite of interest, metabolite regulome networks were constructed using top features identified by Shapley importance analysis (Methods) (Figure 4A). Shapley importance analysis for the top 3 most accurately predicted metabolites for each omics class was conducted to assess the directionality and relative magnitude of feature-metabolite associations (S. Figure 2).

**Figure 4:**
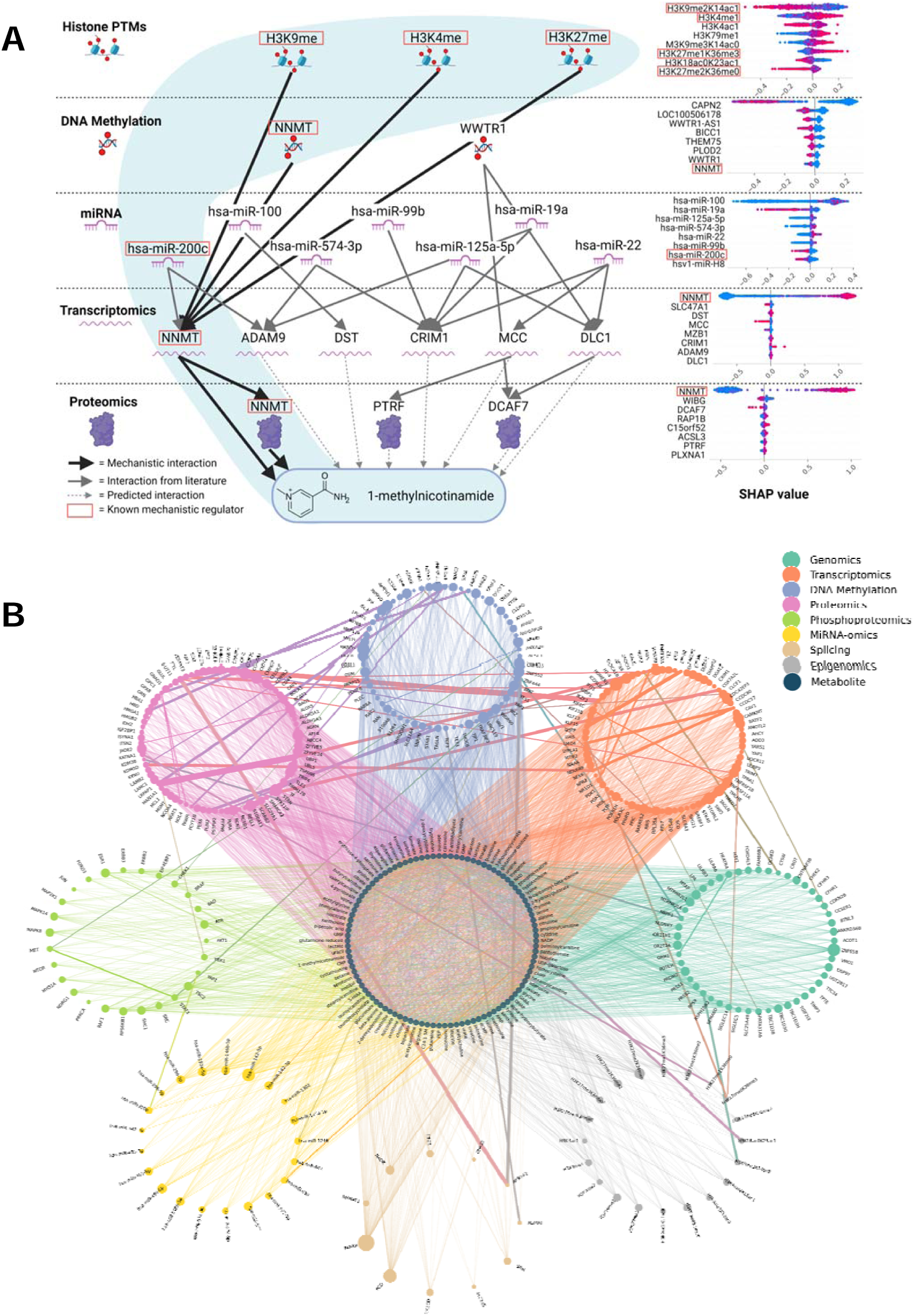
Integrating protein-protein, miRNA-gene and metabolite-gene interactions with Recon8D interactions reveals high consistency between feature importance and topological distribution. **A.** Proposed metabolic regulome network from top 8 features from histone PTM, DNA methylation, miRNA, transcriptomic, and proteomic-trained models for predicting 1-methylnicotinamide (highest mean Shap value) as determined by Shapley analysis. **B.** This figure presents a comparative analysis of relationships between top features derived from Recon8D multi-omics data, including transcriptomics, proteomics, phosphoproteomics, genomics (CNV), methylation, miRNA, splicing and epigenomics, and their corresponding metabolites. Top global features from each omics dataset are organized in distinct, color-coded circular layouts surrounding a central circle of metabolites. The size of each node reflects the number of metabolites associated with that feature. Edges connecting top features to metabolites illustrate the consistency in feature importance and topological distribution across the datasets. Edges between top feature factors represent correlations in their metabolite-consistency profiles, with edge width corresponding to the strength of the correlation. The node selection is based on the top features across the 8 omics datasets. For example, Yap1 shows high consistency with 11 metabolites based on distance of Yap1 to these metabolites in the PPI/miRNA/Recon3D networks and their feature importance in Recon8D, and shares similar metabolite interaction patterns as miR-320b. Met and Stat3 also share similar metabolite interaction patterns with each other and with WIZ (in DNA methylation).

We further focused on 1-methylnicotinamide and phosphocreatine, which were the most highly predictable metabolites (lowest P values and largest correlations) across omics classes (Figure 4A, S. Figure 6). Relationships between miRNA and RNA were identified using TarBase^47^, and between DNA methylation, RNA, and proteins using BioGrid^48^. The interconnectedness of these networks highlights the model’s ability to identify rational biochemical relationships. For example, nicotinamide N-methyltransferase (NNMT), the enzyme involved in the methylation of nicotinamide, was the top predictor of 1-methylnicotinamide levels from both the transcriptomic (CS: 8) and proteomic (CS: 6) models. NNMT has also been shown to modulate histone methylation^49,50^. Similarly, solute carrier family 6 member 8 (SLC6A8) (CS: 8), a creatine transporter, was the top predictor of phosphocreatine in the transcriptomics model, and creatine kinase (CKMT1A) (CS: 3), the enzyme responsible for phosphorylating creatine to create phosphocreatine, was the third most important phosphocreatine predictor in the proteomics model (S. Figure 6). Shapley analysis also identifies the directionality of those relationships. For example, NNMT is positively correlated with 1-methylnicotinamide concentrations (Figure 4A, right). Similarly, hsa-miR-200c, which was identified as a regulator of NNMT translation via TarBase, is negatively correlated with 1-methylnicotinamide levels. The interconnectedness of the top predictive features across models suggests that the relative levels of these metabolites involve high-order interactions between each of these omics layers.

### Physical interactions are frequent among top metabolite predictors

It is difficult to tease out metabolic regulation by individual biomolecules given buffering and combinatorial regulation. By integrating physical and genetic interactions with multiomic associations, we hypothesized that we could understand how groups of regulators complement each other and maintain metabolic homeostasis. We overlaid protein-protein interaction (PPI) data from the STRING database, miRNA-gene interaction data from the TarBase database and metabolite-protein interactions from Recon3D with Recon8D interactions (Figure 4B; S. Figure 5; Methods)^51^. We further analyzed the consistency between the top features in Recon8D with highly connected nodes in the PPI, Recon3D and miRNA networks. We computed a distance matrix between the nodes in the multiple networks and compared it with ML feature importances. Interactions that show high consistency between Recon8D and other interaction networks were visualized (Figure 4B).

This cross-network analysis highlighted several shared interactions between omic layers, especially between various signaling regulators in proteomics and phosphoproteomics, with transcription factors in transcriptomics and DNA methylation. Chek1 in phosphoproteomics shared common patterns of interactions with the transcription factor HOXA5 (DNA methylation); MET and STAT3 (phosphoproteomics) shared similar interactions with each other and with the transcriptional regulators WIZ (in DNA methylation) and the microRNA miR-1246. Similarly, enzymes in fatty acid metabolism shared cross-omic similarity with transcriptional regulators. Lipoic Acid Synthetase (LIAS) enzyme shared similar targets to H3K36 trimethylation, and fatty acid synthase (FASN) with miR-142-3p. Acyl-CoA Synthetase (ACSF3; DNA methylation) had similar targets to aldo/keto reductase AKR7A2 (splicing) with ferritin (FTL), which are all related to redox homeostasis. Genomic copy number changes in Neuroblastoma Breakpoint Family Member 1 (NBPF1) had similar interactions as the GPX8 gene methylation and the histone demethylase KDM3B proteomic levels. (S. Table 13). These results suggest that Recon8D predicted interactions are consistent with known molecular interactions, and our cross-omic analysis highlights potential shared regulation between omic layers.

### Synthetic lethal interactions reveal compensatory control mechanisms

Next, inspired by the synthetic lethal (SL) interaction centered around glutathione highlighted earlier, we systematically assessed for SL interactions across all top metabolite predictors in Recon8D. Since metabolite levels are controlled by multiple factors, we hypothesized that the SL interaction analysis can identify compensatory regulators for each metabolite. We compiled curated experimental human SL interactions from SynLethDB and tested if these interactions are enriched among the top 10 predictors of each metabolite (S. Table 14)^52^. The transcriptome model had the most striking enrichment for SL interactions among top predictors for each metabolite, with 52 interactions involving 45 metabolites (27-fold higher than random chance, p-value = 1 x 10^-34^) (Figure 5A, 5D).

**Figure 5.**
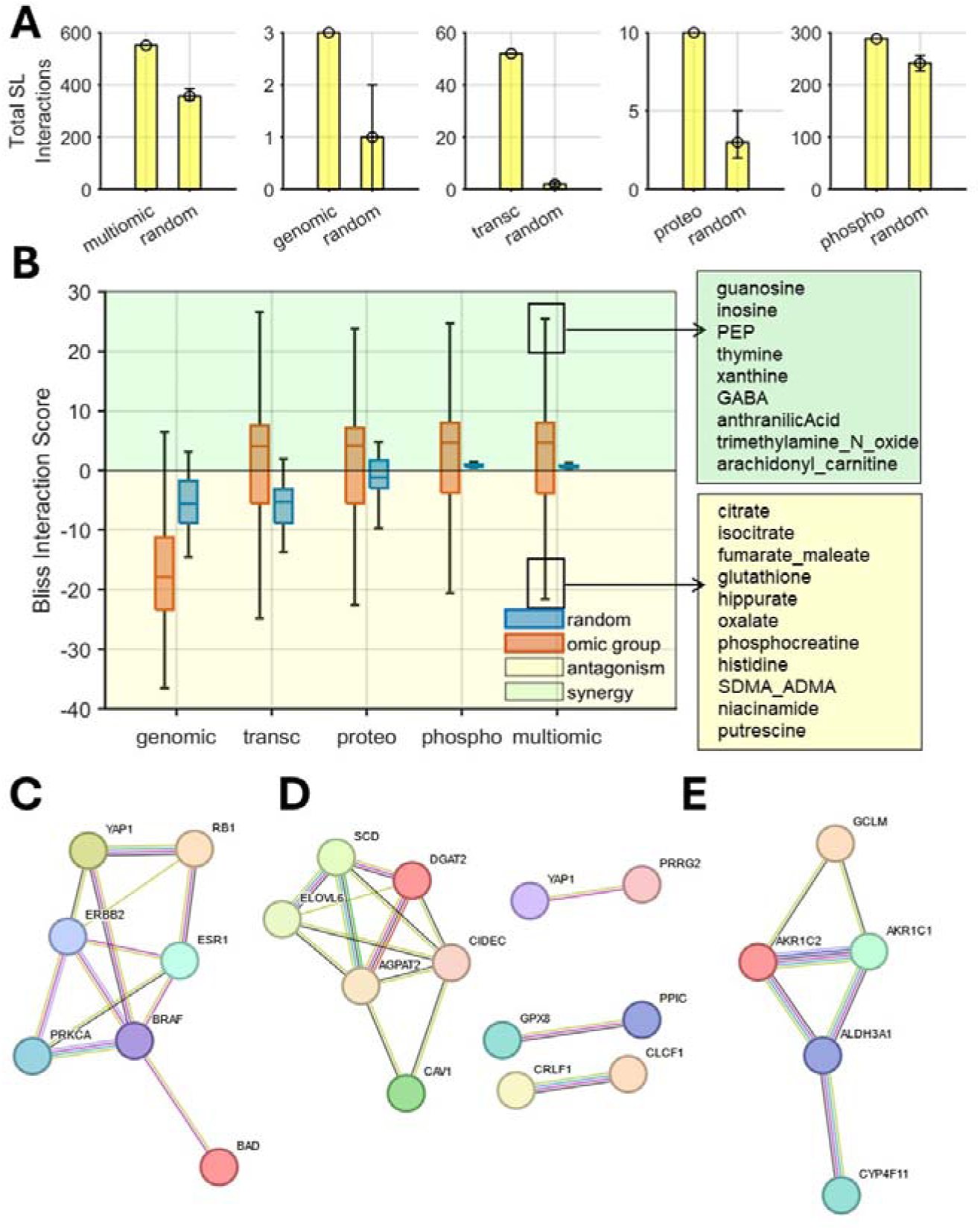
Top metabolite predictors are enriched for synthetic lethality (SL) and drug synergy. **A.** Frequency of SL gene pairs among top 10 influencers using genomic, transcriptomic, proteomic, phosphoproteomic and multiomic (combination of top 10 influencers from genomic, transcriptomic, proteomic, phosphoproteomic ML models). This is compared with frequency of SL pairs obtained from random sampling using the corresponding omics data. SL pairs were obtained from SynLethDB 2.0 which contains 35,943 human SLs. **B**. Average Bliss drug interaction score (positive values implies synergy) for pairs of drugs that target top predictors of a given metabolite derived from various omics. Top metabolites associated with synergy or antagonism are highlighted. Drug interaction data was derived from DrugCombDB. **C**. High confidence genetic and physical interactions among the top 10 global predictors of phosphoproteomics ML model from the STRING database. **D**. High confidence genetic and physical interactions among the top 10 global predictors of transcriptomics ML model from the STRING database. **E**. High-confidence genetic and physical interactions among the top 10 predictors of the metabolite glutathione.

Several SL interactions involve a combination of signaling factors, suggesting that combinations of signaling pathways influence variation in metabolite levels, and each gene in the SL pair may compensate for the other’s absence. For example, the top predictors of nucleotides and nucleotide-precursors showed several SL interactions involving growth signaling factors like AXL, YAP1, CCN1, PTPN14, CSK, and HRAS. Of note, YAP1 and PTPN14 are both top predictors of hypoxanthine, and they both form an SL pair and also physically interact^53^.

SL interactions among top predictors also highlight potential long range regulatory effects. Several interactions involved both a growth signaling and a redox-related gene and were connected to metabolites in central metabolism (Figure 5D, 5E). For example, the top predictive network for lactate contained an SL interaction between the redox enzyme GPX8 and CAV2, which is involved in cell signaling, growth, and lipid metabolism. Similarly, the top predictors for glyceraldehyde-3-phosphate and dihydroxyacetone phosphate, intermediates in glycolysis, also had an SL interaction between GPX8 and the tyrosine kinase AXL.

### Synergistic drug combinations converge on nucleotide metabolism pathways

Synthetic lethality is directly related to drug synergy, wherein drugs targeting proteins in compensatory pathways show enhanced potency when combined. The abundance of SL interactions among top predictors in Recon8D suggests that drugs that target predictors of specific metabolites are likely to be synergistic. To test this hypothesis, we utilized curated drug-drug interaction data in various cancer cell lines from DrugCombDB which includes 448,555 drug combinations covering 2,887 unique drugs and 124 human cancer cell lines^54^. The drugs were mapped onto known targets using data from DrugBank^55^. We tested if synergistic interactions were enriched among drugs that target the top 10 predictors of each metabolite. The transcriptome and phosphoproteome models had the strongest enrichment for synergistic interactions among top predictors for each metabolite (Figure 5B). Drug combinations that target top predictors in the transcriptomic model were highly likely to be synergistic with an average bliss interaction score greater than +4, while randomly selected combinations were likely to be antagonistic (bliss score < -5) (Figure 5B).

Similar to observations with SL interactions, several synergistic interactions involved a combination of signaling pathway inhibitors. We next analyzed metabolites that were associated with synergistic interactions. Notably, nucleotides and nucleotide precursors showed numerous synergistic interactions across a wide range of drugs targeting diverse signaling and metabolic pathways. Guanosine, inosine, pantothenate, thymine, xanthine and hypoxanthine were among the top metabolites associated with most synergistic drug combinations (Figure 5B, S. Table 15). Out of 104 drug combinations associated with these metabolites, 100 were synergistic (S. Table 15). Drugs that target diverse signaling pathways may lead to similar downstream effects on nucleotide metabolism. For example, several tyrosine kinase inhibitors including sorafenib and sunitinib are predicted to be associated with these nucleotides (S. Table 15). These drugs are known to affect levels of nucleotides including guanine, adenine, adenosine, guanosine, hypoxanthine, and inosine^56^. To further validate these associations, we used the DepMap Data Explorer to quantify the relationships between cellular sensitivity to these signaling drugs and levels of nucleotides, and found several statistically significant associations (S. Table 16)^46^. These results confirm the connection predicted by our analysis. In contrast, antagonistic drug interactions were surprisingly associated with metabolites in the TCA cycle. Citrate, isocitrate, and fumarate were the top metabolites associated with most antagonistic drug combinations. Several drug combinations associated with these TCA cycle metabolites involve drugs such as Sirolimus, Everolimus and Tetracycline that are known to reduce the activity of the TCA cycle (S. Table 15)^57,58^. Drugs that perturb TCA cycle activity may lead to antagonism with agents that require TCA cycle activity. Thus, Recon8D can provide insights into understanding combinatorial metabolic regulation, synthetic lethality, and drug synergy.

## Conclusion

Cellular metabolism is controlled by multiple layers of regulatory networks. Here, we quantified the influence of potential regulatory interactors on metabolomic variation from matched omics across 8 biomolecular classes. The metabolome was surprisingly highly predictable from transcriptomic data. While the absolute levels of metabolites may depend on thermodynamics, variation may result from differences in transcriptional regulation. Further, the relationship between transcripts and metabolites is nonlinear, and a combination of transcripts influences metabolite levels. In many cases, there is no direct correspondence between metabolites and enzymes they are involved with. Signaling-related transcripts, proteins, and phosphoproteins were top predictors across omics layers. A second common global predictor was redox metabolism, exemplified by GPX enzymes, that influence glutathione levels and overall redox state of the cell. Levels of histone PTMs, miRNA and phosphoproteins were also highly predictive of the metabolome despite having the fewest features by an order of magnitude. Analysis of feature importance levels of individual metabolites revealed that metabolites in peripheral metabolism, like taurine, were most predictable via levels of enzymes related to those metabolites, while metabolites in central metabolism were more predictable by a combination of signaling and redox regulators. Because central carbon metabolism has been more extensively studied than peripheral metabolism, we would expect to find strong clear associations with central carbon metabolism. However, we find the opposite, that central carbon metabolites often do not have a clear top predictor, suggesting that these metabolites are more prone to combinatorial regulation. We found these patterns to hold true in independent datasets from NCI60 and single-cell omics. Our analysis thus uncovered key systems-level insights on metabolome variation between cells, the generalizability of ML models, and predictability of individual metabolites.

Although we make use of ML to predict the metabolome, such models are not directly transferrable in predicting metabolomics from other sources due to several factors, including differences between *in vitro* and *in vivo* conditions, and between cell line panels, and limited features overlap within and across omics. Furthermore, one of the key observations from this study is related to the predictability of specific metabolites, such as those in central metabolism, which can affect the generalizability of ML models that aim to predict metabolomes. While many metabolites were predicted in our Recon8D model to be preferentially influenced by a single top feature, others were predicted by multiple features, suggesting that they may be regulated combinatorially. Such metabolites will be harder to predict using a single omics feature. Hence, the focus of Recon8D is not the prediction of absolute metabolite levels, but rather the associations between features and metabolites, as reflected in our validation experiments.

A limitation of our study is that our analysis was done in cancer cell lines in culture and may not reflect metabolism *in vivo*; however, patterns observed here related to transcriptional variation, and topological relatedness of gene-metabolite associations have also been observed *in vivo* in humans and model systems^39,59,60^. Further, the ’guilt by association’ strategy used by network inference methods may not identify causative mechanisms. We address this through integration of multiple layers of orthogonal omics information that we found to be consistent with each other^61^. Further, these interactions are robust across hundreds of cell lines in independent datasets spanning multiple tissue lineages and growth conditions. Our model also correctly recalls several known interactions reported in literature. The above suggests that the statistically inferred associations may represent real interactions.

Another limitation is that network associations driven by systems-level data are descriptive, and findings are not necessarily mechanistic, even among high confidence features. However, along with extensive documentation of Recon8D associations reflected in known molecular interactors, these findings suggest strong mathematical relationships between the metabolome and other omics classes. Experimentally validating proposed interactions from this study, even for a small percentage of interactions, would require an immense effort. We hope that these prioritized associations may inform rational exploration of novel metabolic regulatory interactors.

Additionally, some high-throughput measurements do not provide a comprehensive panel of molecules that exist within a cell. For example, the metabolite panel used for this study consisted of 225 metabolites, a fraction of those that exist within a cell across distinct cellular compartments. Furthermore, there exist more proteins in human cells than used in this study, and the 90 phosphoproteins present were measured using a targeted approach, and therefore are not representative of the entire cellular array of phosphoproteins. To make a strong inference as to the relative influence of these classes on the metabolome, these analyses would need to be conducted on more comprehensive, untargeted data. Similarly, differences in sample quantities can influence accuracies, where omics with too few samples (cell lines) may not perform as well (S. Figure 1, S. Table 1).

Beyond the relative predictability of the metabolome from omics inputs, we find that top predictors from ML are mechanistically related to the associated metabolites. For example, in the transcriptomics ML model using genes present in Recon3D, the metabolites’ top predictors were enzymes closely associated, either directly (17%) or first-degree neighbor (81%), with those metabolites. The top predictors for each metabolite across different omics layers were highly enriched for genetic and pharmacological interactions. This further supports the ability of our approach to uncover mechanistic interactions and highlights potential multiomic compensatory buffering between top metabolomic predictors. Finally, we find that drugs that target top predictors in nucleotide metabolism are likely to be more synergistic. This builds upon our observation of increased synthetic lethality among the top predictors of each metabolite. Notably, a wide range of signaling drugs intersect with nucleotide metabolism and are predominantly synergistic when used in combination (mean bliss score > 10, S. Table 16, Figure 5B). Our analysis suggests that these drugs, which target signaling pathways, may also impact nucleotide homeostasis. This is consistent with a prior study that found combinations of Mek and PI3 Kinase inhibitors show synergy via nucleotide metabolism^62^. In contrast, drug antagonism was associated with metabolites in the TCA cycle. A similar observation was made in bacteria wherein antibiotics that have opposing effects on TCA cycle and bacterial respiration are antagonistic^63^. Several drugs require active growth and metabolism to be potent. Inhibition of TCA cycle may result in growth arrest which may lead to reduced potency of growth-dependent drugs. In contrast, cells with nucleotide imbalances continue to grow^64^. This may result in errors in DNA replication and transcription leading to cell death. Overall, Recon8D may provide a rational framework for metabolic engineering and designing synergistic combination therapies.

## Methods

### Data Acquisition and Preprocessing

Omics from the Cancer Cell Line Encyclopedia (CCLE) were obtained from the Broad Institute (Cambridge, MA) Dependency Map Portal (with the exception of proteomics and phosphoproteomics, which were obtained from Nusinow et al and Li et al, respectively)^28,65,66^. These include genomics (copy number variation and mutations), epigenomics (bulk histone H3 modifications (MRM) and DNA methylation (RRBS)), transcriptomics (RNAseq), RNA splicing (RNAseq, exon inclusion ratio), miRNA-omics (expression), lncRNA (expression), proteomics (mass spectrometry), phosphoproteomics (RPPA), and metabolomics (LC-MS). The number of cell lines used for training can be found in S. Table 1.

Data preprocessing involved replacing missing values (NaN) with zeros, taking the median value of repeated observations (if required), standardizing cell line or feature names across datasets, and removing features with 0 variance. Only lncRNA, RNA splicing, and mutation data had features with zero variance (38, 1,131 and 146 features, respectively). Once removed, there were 3,767, 2,957 and 17,203 features used for model training, respectively. RNA splicing ensemble IDs were converted to common names using BioTools.fr^67^. Each data type was merged independently with the metabolomics dataset by common cell line, where each matrix was organized by cell lines (rows) and features (columns). Input datasets for the models containing metabolically relevant genes were filtered for genes present in the Recon3D genome-scale metabolic model^68^. The same data preprocessing steps and scaling methods were used for validation experiments with NCI60 data, which were obtained from the National Cancer Institute CellMiner portal^29,69^.

### Machine Learning Model Training

ML models that predict metabolomic variation were trained on individual omic data from the CCLE. A separate model was trained and tested for each metabolite using 5-fold cross-validation, where data were normalized by Z-score within each CV fold. Statistically significant predictions were determined using a Bonferroni-corrected P value (0.05/225) to account for false discovery rate. Because the number of features was much greater than the number of observations in these data, it was important to select algorithms that penalize large coefficients and limit overfitting. Random forests were primarily used for this study, although several algorithms were tested for benchmarking, including Ridge Regression, Lasso Regression, XGBoost, and SVMs^70–74^. Each of these algorithms is easily interpretable (provides feature importance scores or coefficients) and characterizes different mathematical relationships. Non-linear SVMs are more difficult to interpret, and therefore mutual information calculations were required to provide surrogate feature importance scores. The Python script for these algorithms can be found at the Recon8D <u>GitHub repository</u>. The trained random forest models, along with the CCLE data used for model training, can be found on the <u>Recon8D Synapse</u> page.

Random forests are comprised of many decision trees, and are classified as an ensemble learning method, where they aggregate many iterations of decisions. This algorithm works by randomly selecting a training set from the input data (bootstrapping) and using that training set to dictate a unique decision tree^75^. These predictions are averaged over all the decision trees, which is what allows for more accurate predictions from the overall model than any individual tree could provide^75^. Ultimately, we chose random forests for the bulk of our analyses for their balanced and interpretable framework as well as their overall high accuracy in comparison to other algorithms. More complex models can be challenging to interpret, and simpler models may not identify non-linear or combinatorial relationships.

### Hyperparameters

Random forests were trained using Sci-kit Learn’s Random Forest Regressor algorithm with the following parameters: (random_state=0, n_estimators=100, n_jobs=-1, max_depth=10). These parameters were selected after considering computation time while maintaining sufficient depth. While random forests are robust to changes in hyperparameters, we elected to not tune such parameters to avoid overfitting. A similar approach was taken for XGBoost and SVMs.

For ridge and lasso regression, an array of alphas was used in conjunction with cross validation to determine the appropriate regularization term for each model. We used an array containing alpha values of 1e-2, 0.1, 1, 10, 100, and 1000. The best alphas were saved for each cross-validated model and applied to final model training using the whole data.

### Model Evaluation and Interpretation

Performance of each model was assessed using Pearson’s correlation between predicted and true metabolite concentrations. FDR corrected P values were also calculated for assessing significance. Cross-validation was used to assess overall accuracy of each data type, and a final model trained on the entire data set was used to examine feature importances. Feature importance scores were generated from each random forest model and were analyzed locally (for each metabolite) and globally (averaged across all metabolite models for a given omics type). Global importances were sorted by median rank and scaled by Z-score (S. Table 10).

Mutation feature importances for SVMs were calculated using mutual information—a model-agnostic method for assessing the dependency between a feature and a metabolite. This was done in lieu of directly interpretable model characteristics of non-linear SVMs. Instances of mutations in the top 20 features for each metabolite (similar to control experiment confidence scores) were added to assess the impact of each mutation in predicting the metabolome.

Feature importances for select predicted metabolites were also quantified using Shapley analysis, a game-theoretic approach that calculates feature importances by adding each feature into the model sequentially, and repeating that process for all orderings of features, thereby assessing the influence of each feature on the model as a whole^76,77^. The three metabolites with the lowest P values were selected for Shapley analysis for each data type (S. Figure 2).

Consistencies across control experiments were quantified with a confidence scoring system, where the top 20 features were extracted from whole-data and control models. We then tallied the occurrence of each of the top 20 features conserved in the top 20 features of each control experiment, out of 8 experiments, leading to a confidence scoring system ranging from 0 to 8 for each feature. The 8 experiments included models generated on the following subsets: lung cancers, hematopoietic and lymphoid cancers, all cancers except lung, all cancers except hematopoietic and lymphoid, suspension cell culture, adherent cell culture, primary tumor samples, and metastatic samples.

The top one, two, five and ten features (quantified by random forest feature importance scores) from genomics (CNV), DNA methylation, transcriptomics and proteomics models (for metabolic gene-only models) were mapped back to the Recon3D metabolic model reactions. This was accomplished by matching metabolites from the CCLE panel to metabolites present in the Recon3D metabolic model. Common metabolic cofactors were removed to avoid overrepresenting neighboring reactions, including ATP, ADP, water, hydrogen, phosphate, and diphosphate, which may be found in the recon_mapping section of the <u>GitHub repository</u>. Functions from the COBRA toolbox allowed for the identification of all reactions associated with the relevant metabolites (both primary and first-degree neighbors), as well as the genes associated with each reaction^78^. Once identified, gene names were converted to BiggIDs and matched to each set of reactions. The number of metabolite models that had at least one top feature present in the relevant set of reactions (out of the 95 metabolites in both the CCLE panel and Recon3D) were compiled (S. Table 9). We performed this same analysis on a set of random transcripts, which show fewer overlaps (S. Table 9). A random set of 5 transcripts was taken for each metabolite and overlaps were counted. This process was repeated 50 times, and the median value of overlaps is represented in the table.

### Model Validation in independent datasets

The top features from CCLE-trained models were isolated and used to analyze relationships between transcripts and metabolites in the NCI60 data and single-cell data. Random forest feature importance scores from NCI60-trained and single-cell-trained models were used in comparison to CCLE outputs, where top features as identified by CCLE models were isolated in NCI60 data to examine if they were larger than by random chance. Z-score-scaled feature importance ranks were used for this analysis. Student’s t tests and Kolmogrov-Smirnov tests compared isolated feature importances to the entire feature set. Data were stacked to create one distribution among all metabolites (pooled). Aggregated importances showed significant differences between the top features and overall feature set. These analyses were conducted on sets of 1, 3, 5, 10, 50, and 100 top features from each metabolite model (except for phosphoproteomics, which only had 90 features, and thus the 100-feature subset was not used).

### Effect of feature size, missing features and metabolic features

We explored how well metabolic genes alone could predict the metabolome. Genomic, DNA methylation, transcriptomic, and proteomic datasets were filtered for genes present in the Recon3D metabolic model^68^, and new models were trained on the metabolic gene subset. The difference in feature size between omics classes can affect their predictive power, and filtering for genes present in Recon3D standardized these data to provide mechanistic information within the context of known metabolic networks. Each filtered dataset had a similar number of features, which provided an avenue through which to assess the impact of feature size on model accuracy.

We first filtered the feature sets for metabolically relevant genes from Recon3D (S. Figure 4A). With fewer than 2,000 genes, transcriptomics inputs predicted over 99% of metabolites statistically significantly, and other feature sets conferred similar accuracy to whole-data models in predicting each of the 225 metabolites (S. Figure 4B). The median correlation for each omics class was slightly lower in the metabolic gene-only models as compared to the whole-data models, but the distributions were similar overall (S. Figure 4C). Similar accuracies from the metabolic gene-only models and the whole-data models illustrate the lack of impact of feature size on model predictions. We further explored this by training models on a random set of 2,000 genes, which also had similar accuracies. While models built on randomly selected features might not provide mechanistic insights, they show that the metabolome is highly predictable within a reasonable range of input feature sizes. Feature importances from metabolic gene-only models and whole-data models were also conserved significantly (S. Figure 4D and E).

Both the number of training samples (cell lines) and features can impact accuracy. Proteomics and phosphoproteomics had fewer cell lines for training than the rest of the omics (S. Table 1), which could explain lower accuracies comparatively. Additionally, we hypothesize that the reduced feature size in the proteomic data may account for the reduced accuracy in these models compared to transcriptomics. We interrogated this by comparing the transcriptomic and proteomic datasets to see if there were discrepancies in the feature set that might influence accuracy. Between the 416 genes that were present in the transcriptomics feature set but were absent from proteomics, 11 of those genes were the top 1 predictor in a metabolite model. These missing features explain the discrepancy in accuracy (∼15%) between transcriptomic and proteomic inputs, suggesting that access to more comprehensive proteomic data might be more predictive of relative metabolite levels.

### Statistical analysis

All P values were corrected using Bonferroni multiple hypothesis correction. Statistical significance of the overlaps with synthetic lethal interactions from SynLethDB 2.0 and drug interactions from DrugCombDB were assessed by creating 50 sets of random top features for each metabolite in all omics models and quantifying the overlap. This was then compared with the actual overlap using a t-test.

### Multi-Omics Topological Analysis Based on Metabolic, miRNA and PPI Networks

The metabolic network data was obtained from the Recon3D genome-scale metabolic model. Protein-protein interaction (PPI) data was sourced from the STRING database (https://string-db.org/), and miRNA-gene interaction data was sourced from the TarBase database (https://dianalab.e-ce.uth.gr/tarbasev9). A metabolic network linking metabolites to genes was constructed based on the metabolite-reaction-gene relationships provided by the Recon3D genome-scale metabolic model. The PPI network and miRNA-gene network were constructed directly using the pairwise relationships extracted from the STRING and TarBase databases, respectively. The Construction of Distance Matrices and Rank Difference Analysis involves two steps. First, the top features (top 20 for epigenomics, top 40 for phosphoproteomics, top 100 for other omics) were selected by feature importance ranking, mapped to the PPI network, and used to compute an n×l distance matrix (n: number of top genes in the PPI network; l: number of shared genes/proteins between the metabolic and PPI networks). Next, an l×m metabolic distance matrix was generated from the positions of genes in the Recon3D metabolic network (m: metabolites common to the metabolic network and CCLE metabolomics data). The final n×m omics-metabolite distance matrix was obtained by multiplying these matrices. For miRNA, a three-step process was used: first, a distance matrix based on miRNA-gene interactions was generated, followed by multiplication with the previous two matrices to yield the final omics-metabolite distance matrix. To compare the ranking differences of top features between the distance and feature importance matrices, both were converted into ranks, and the absolute differences were normalized to a range of 0 to 1. For each metabolite, the top 10% and bottom 10% of feature-metabolite relationships were classified as high and low consistency, respectively. Cross-omic edges were determined based on similarity of metabolite interaction patterns based on the Pearson’s correlation coefficient, and a threshold of +0.4 was used to visualize significant cross-omic similarities.

## Acknowledgments

This work was supported by faculty start-up funds from the University of Michigan (UM), Camille and Henry Dreyfus Foundation, the Rogel Cancer Center at UM, R35 GM13779501 from NIH to SC, and the Advanced Proteogenomics of Cancer (T32 CA140044) Program to RS. We thank Deepak Nagrath, Maciek Antoniewicz, Analisa Difeo and Yatrik Shah for suggestions and critical feedback on the project.

## Competing interests

The authors declare no competing interests.

## Data availability

All datasets are provided in figures, the supplementary materials, and in the GitHub repository.

## Author contributions

S.C, R.S designed and performed research, and wrote the manuscript; K.S, R.B, Y.L, S.M, A.C.V, E.K, M.K, S.N performed research.

## Supplementary Materials

**Supplemental Figure 1:**
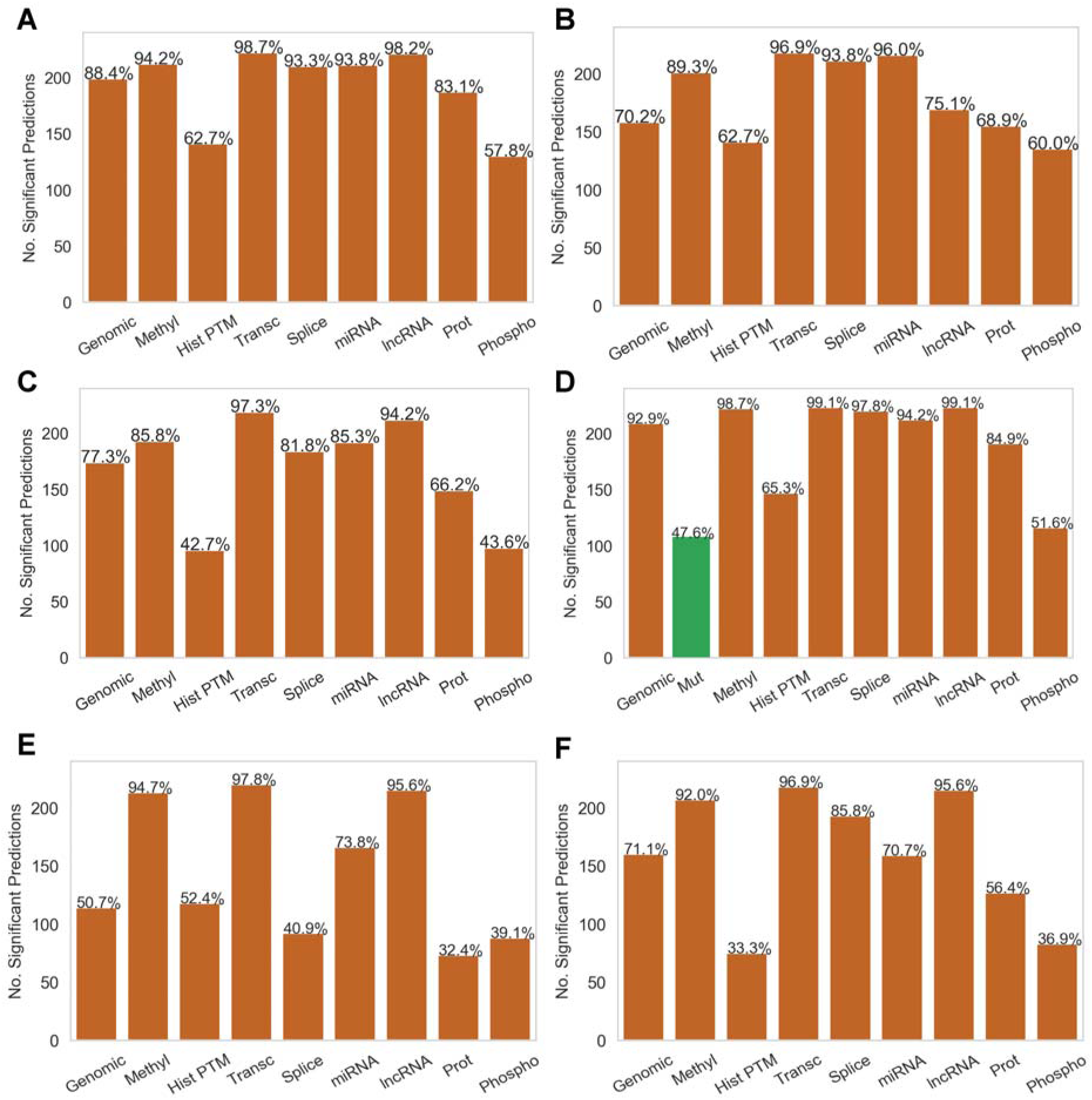
Results from training an array of ML models on each input dataset. Bar charts show the number of significantly predicted (FDR p-value < 0.05) metabolites (of 225) per input data type, also represented as a percentage out of 225. These algorithms did not significantly outperform Random forests, and therefore were not selected for in-depth analysis. **A.** Random Forest results for comparison (also shown in Figure 2) **B.** Results from Random Forest benchmarking experiments using the top 40 features per input dataset as determined using 75-25% holdout validation. **C.** XGBoost **D.** Support Vector Machine (SVM) with RBF kernel. Highlighted in green are results from binary mutation data **E.** Ridge regression **F.** Lasso regression

**Supplemental Figure 2:**
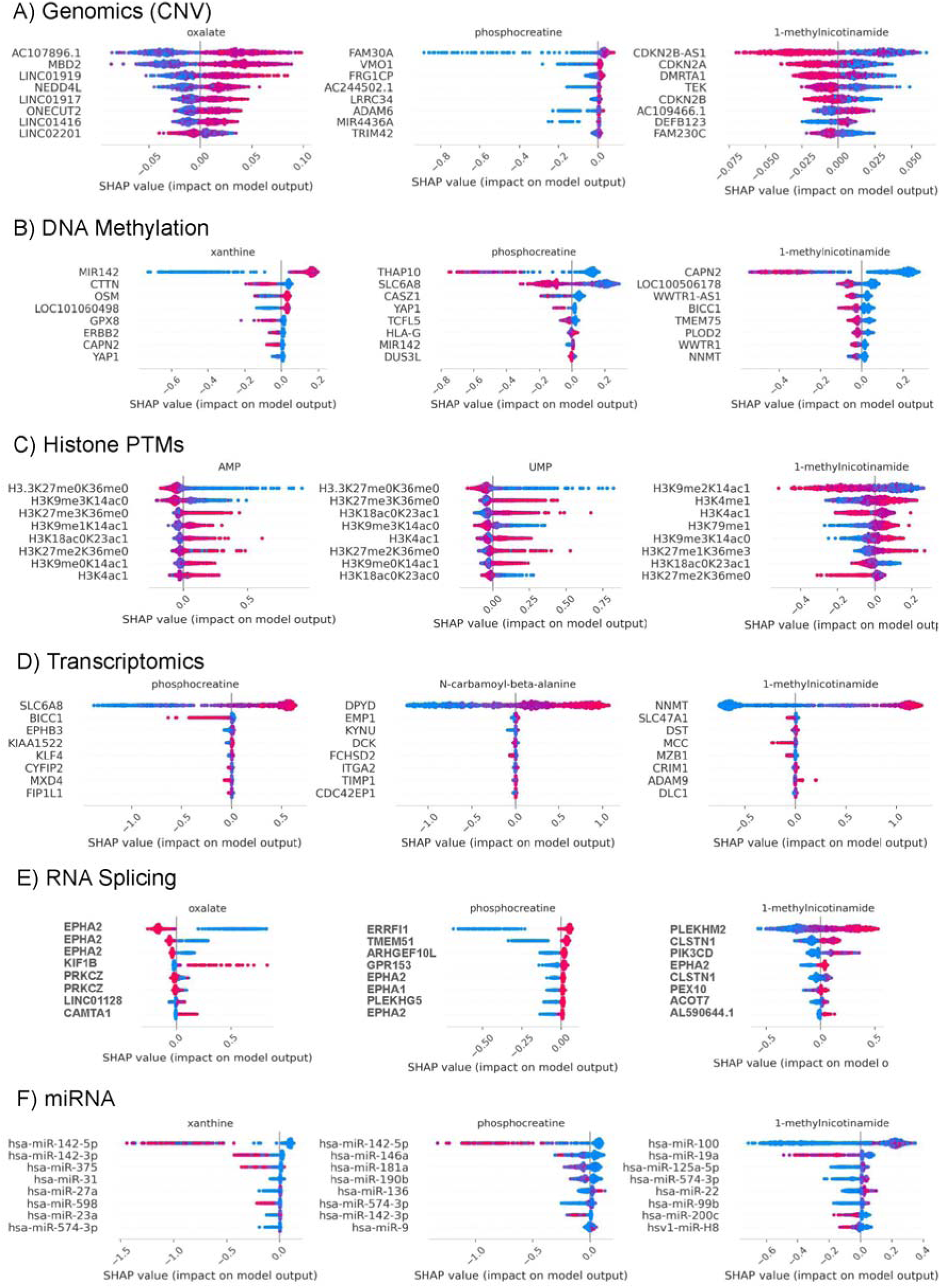
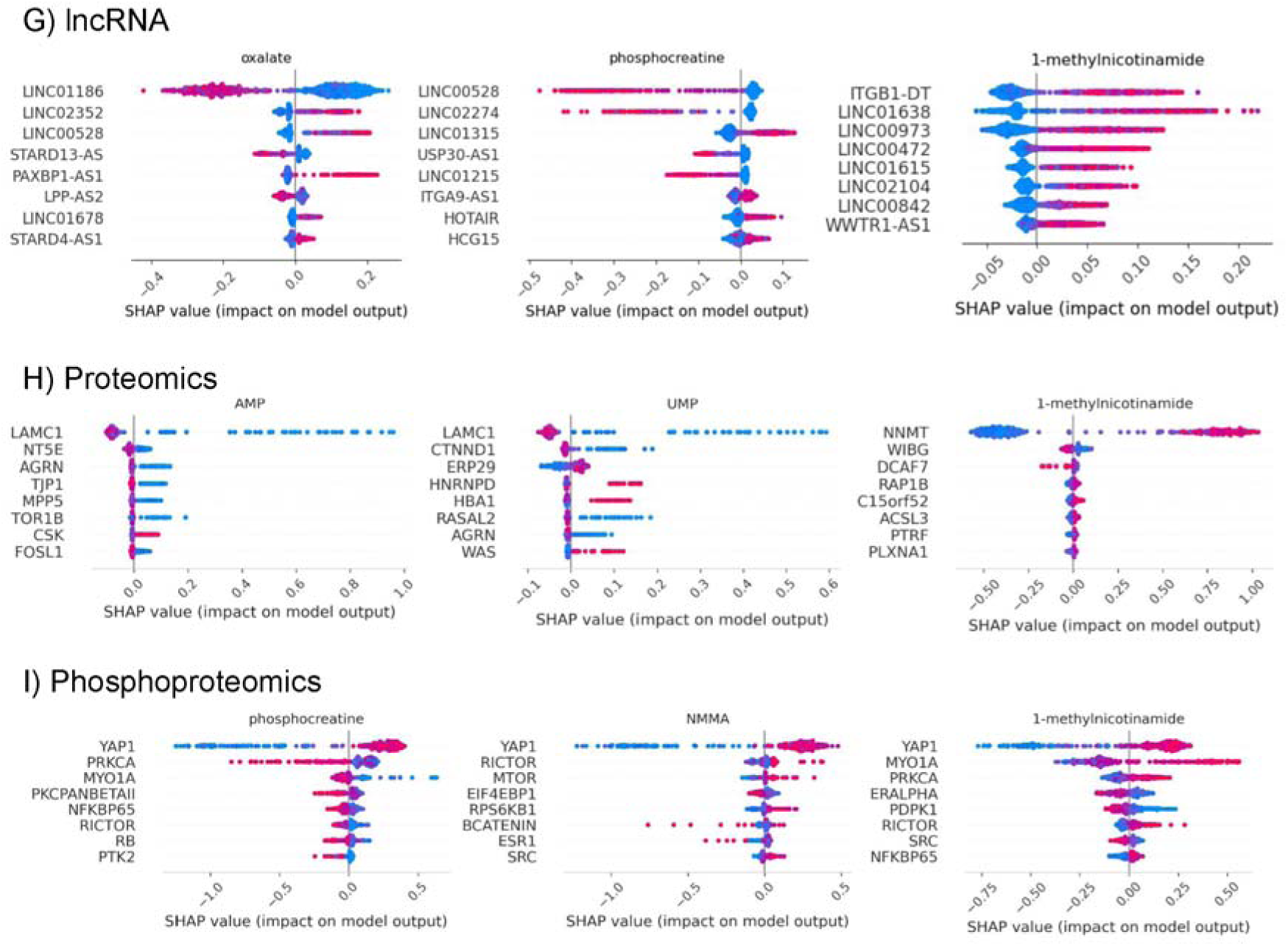
Shapley analysis for top 3 metabolites (lowest P value) for each omics class from random forest models. Each dot on the Shapley beeswarm plots corresponds to a cell line observation. Red data points indicate high feature values and blue points indicate low feature values. High SHAP values correspond with higher metabolic concentration outputs from the model, and low SHAP values correspond with lower outputs. Therefore, a gene with a high feature value and SHAP value (red and to the right) illustrates a positive correlation, and a high feature value and a low SHAP value (red and to the left) illustrates a negative correlation. **A.** Genomics (CNV) **B.** DNA methylation **C.** Histone PTMs **D.** Transcriptomics **E.** RNA Splicing **F.** miRNA **G.** lncRNA **H.** Proteomics **I.** Phosphoproteomics

**Supplemental Figure 3:**
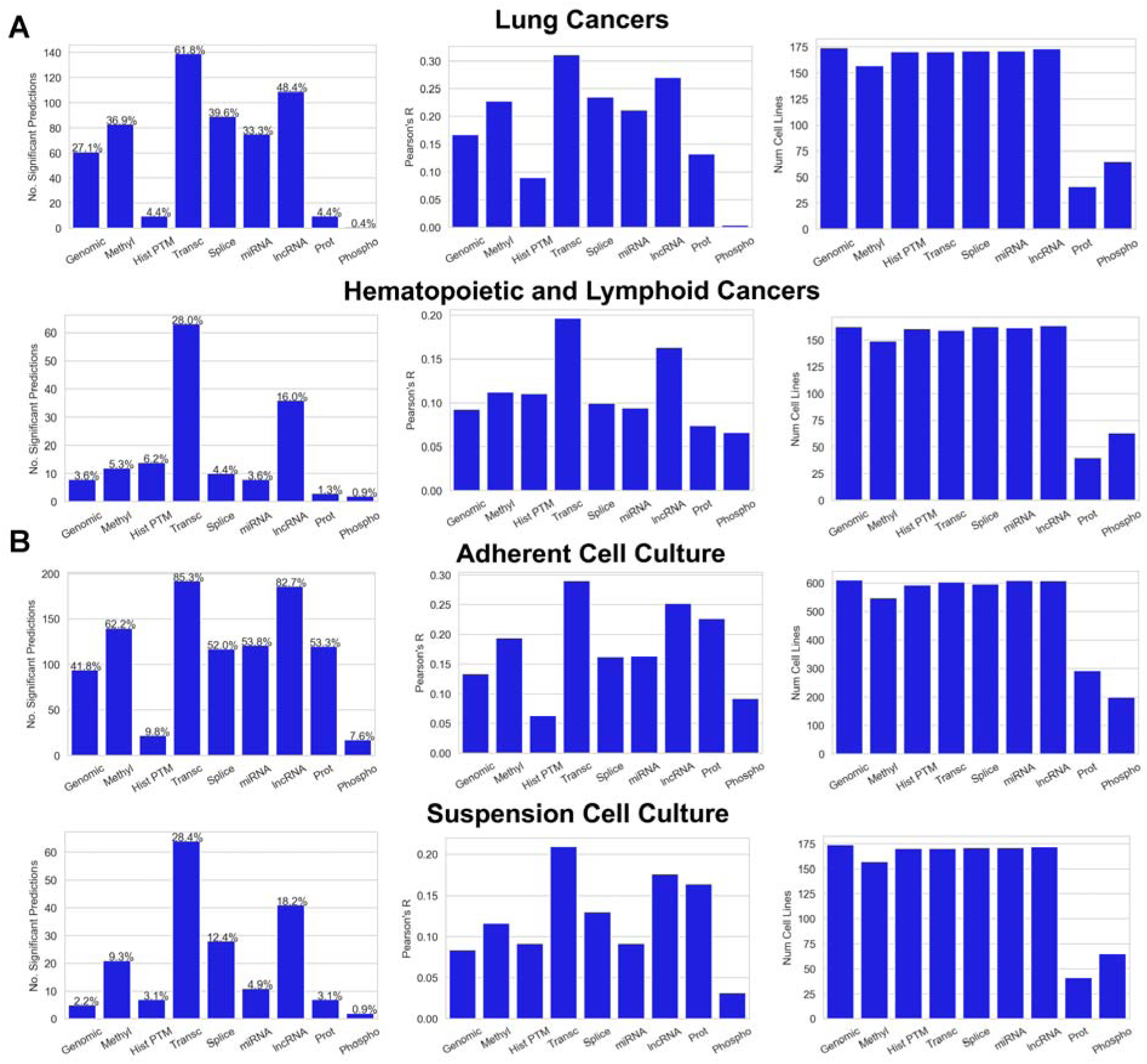
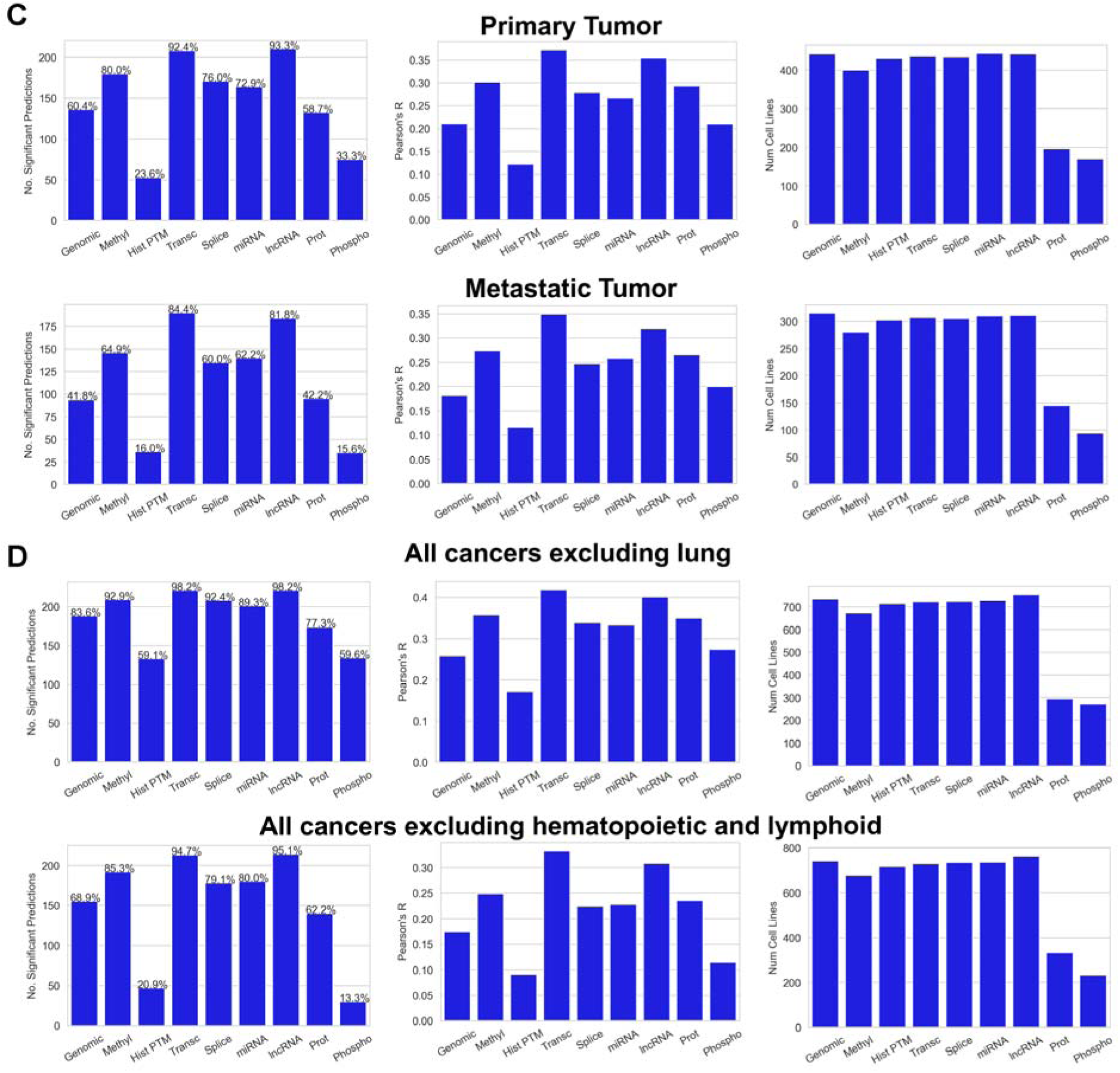
Control experiment results (random forests), showing the number of significant predictions (FDR p-value < 0.05) out of 225 metabolite models (also represented as a percentage out of 225), the average correlations across all models, and the number of cell lines in each data set (left to right). **A.** Models trained on samples from particular cancer lineages. **B.** Models trained on samples grouped by culture conditions. **C.** Models trained on samples grouped by tumor origin. **D.** Models trained on all cancer lineages excluding lung cancers or excluding hematopoietic and lymphoid cancers.

**Supplemental Figure 4:**
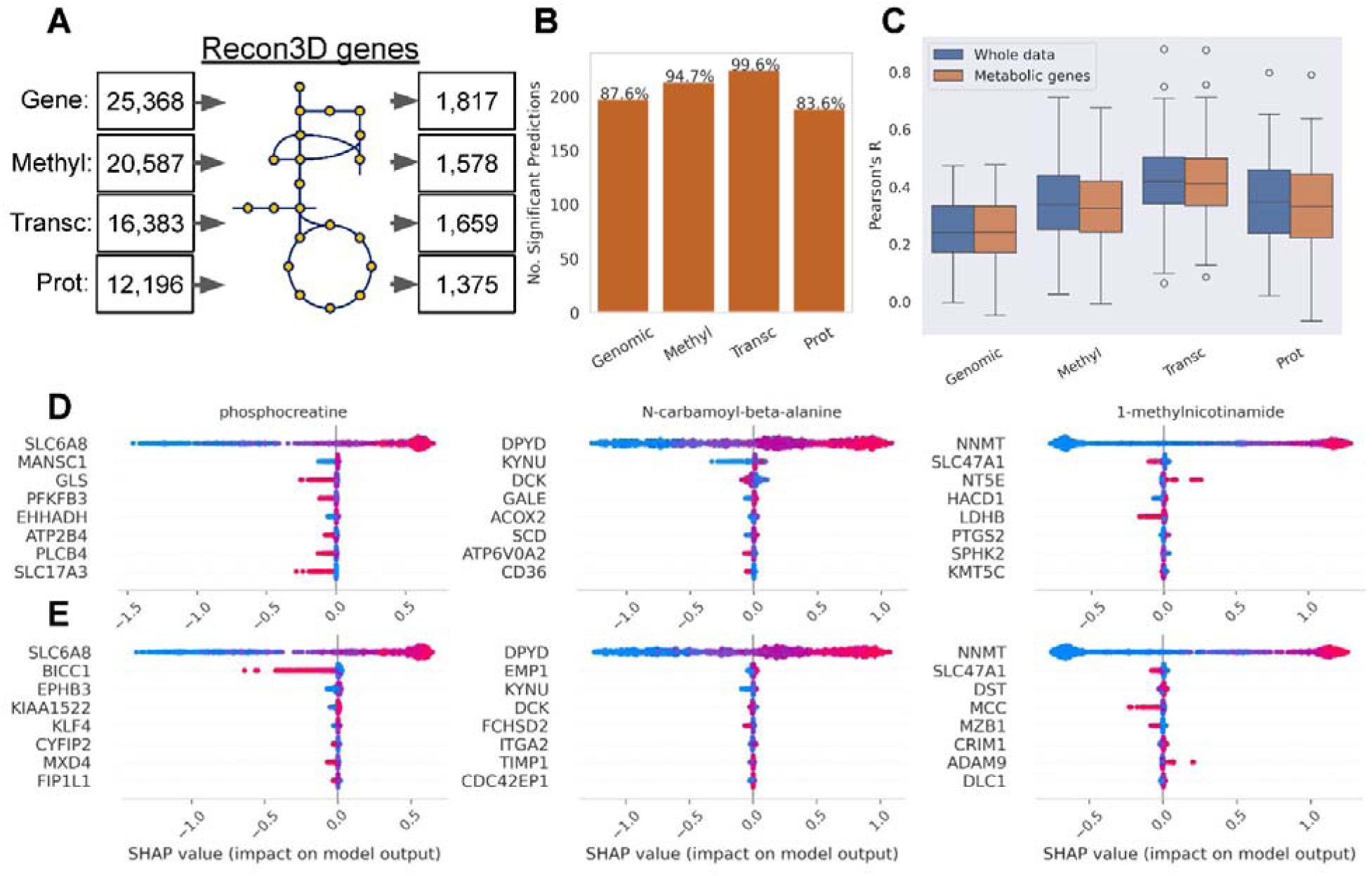
Results from models trained on genes present in the Recon3D metabolic model. **A.** Number of features in each omics class before and after filtering for genes present in the Recon3D metabolic model. **B.** Number of significant predictions (FDR p-value < 0.05) from metabolic gene-only trained models, also represented as a percentage out of 225. **C.** Box plots showing distribution of correlations (true vs predicted) from models built on whole data versus metabolic gene-only models (right). **D.** Shapley analysis for top 3 metabolites (lowest P value) from metabolic gene-only models identifying the top 8 most important features. Each dot on the Shapley beeswarm plots corresponds to a cell line observation. Red data points indicate high feature values and blue points indicate low feature values. High SHAP values correspond with higher metabolic concentration outputs from the model, and low SHAP values correspond with lower outputs. Therefore, a gene with a high feature value and SHAP value (red and to the right) illustrates a positive correlation, and a high feature value and a low SHAP value (red and to the left) illustrates a negative correlation. **E.** Shapley analysis for top 3 metabolites (lowest P value) from models built on whole data identifying the top 8 most important features. These show that feature importances from metabolic gene-only models and whole-data models were conserved. For example, NNMT is the top predictor for 1-methylnicotinamide, SLC6A8 for phosphocreatine, and DPYD for N-carbamoyl-beta-alanine in both the metabolic gene-only and whole-data models. DPYD is involved in N-carbamoyl-beta-alanine biosynthesis.

**Supplemental Figure 5:**
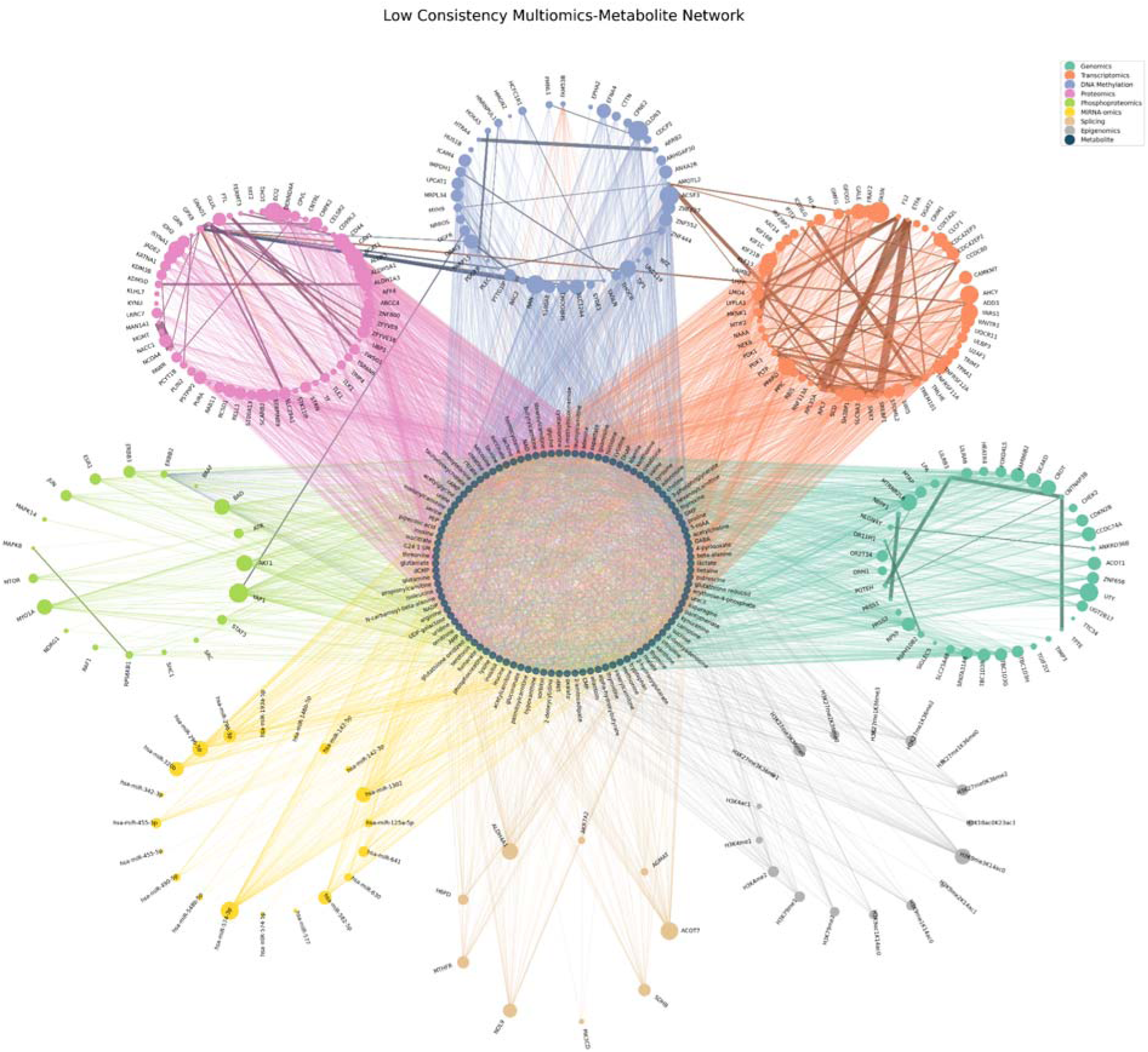
Integrating protein-protein, miRNA-gene and metabolite-gene interactions with Recon8D nteractions. This figure presents a comparative analysis of relationships between top feature factors derived from Recon8D multi-omics data, including transcriptomics, proteomics, phosphoproteomics, genomics, methylation, miRNA, splicing and epigenomics, and their corresponding metabolites. Top features from each omics dataset are rganized in distinct, color-coded circular layouts surrounding a central circle of metabolites. The size of each node eflects the number of metabolites associated with that feature. Edges between top features represent correlations in heir metabolite-consistency profiles, with edge width corresponding to the strength of the correlation. This figure resents a comparative analysis of low consistency relationships between top feature factors derived from multi-mics data. Edges connecting top feature factors to metabolites illustrate the variability (in contrast to Figure 5, which shows consistency) in feature importance and topological distribution across the datasets. The figure emonstrates how feature importance and topological distribution can diverge (low consistency) across multi-mics data. For example, Yap1 (phosphoproteomics) shows low consistency with 60 metabolites based on distance f Yap1 to these metabolites in the PPI/miRNA/Recon3D networks and their feature importance in Recon8D.

**Supplemental Figure 6:**
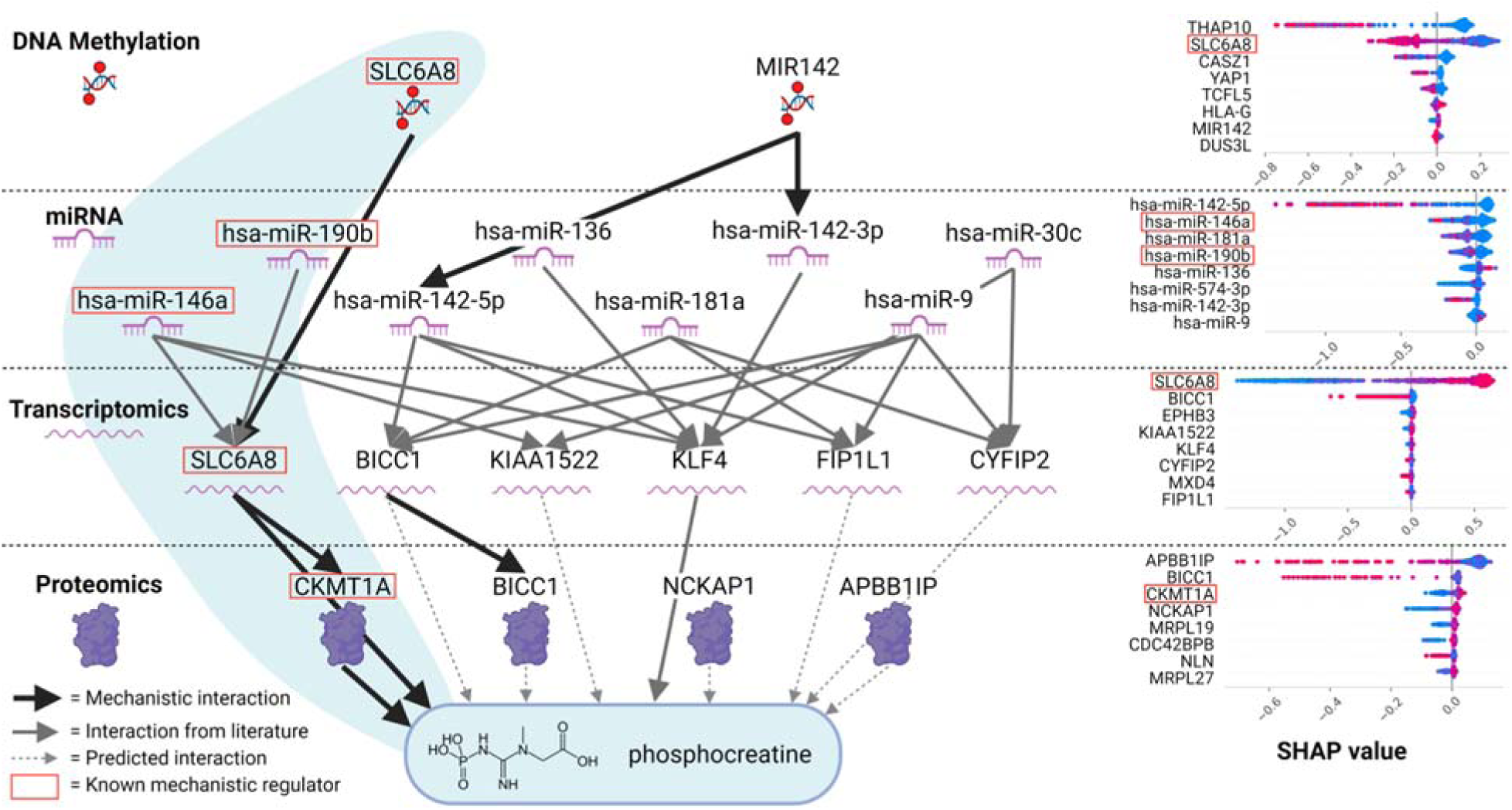
Proposed metabolic regulome network from top 8 features from miRNA, transcriptomic, and proteomic-trained models for predicting phosphocreatine (highest mean Shap values) as determined by Shapley analysis.

**Supplemental Table 1:** Number of cell lines used to train each model.

| Cell Lines | Genomics | Mutations | DNA methylation | Histone PTMs | Transcript-omics | RNA Splicing | miRNA | lncRNA | Proteomics | Phospho-proteomics |
| --- | --- | --- | --- | --- | --- | --- | --- | --- | --- | --- |
|  | 921 | 925 | 828 | 893 | 904 | 900 | 916 | 916 | 373 | 296 |

**Supplemental Table 2:**
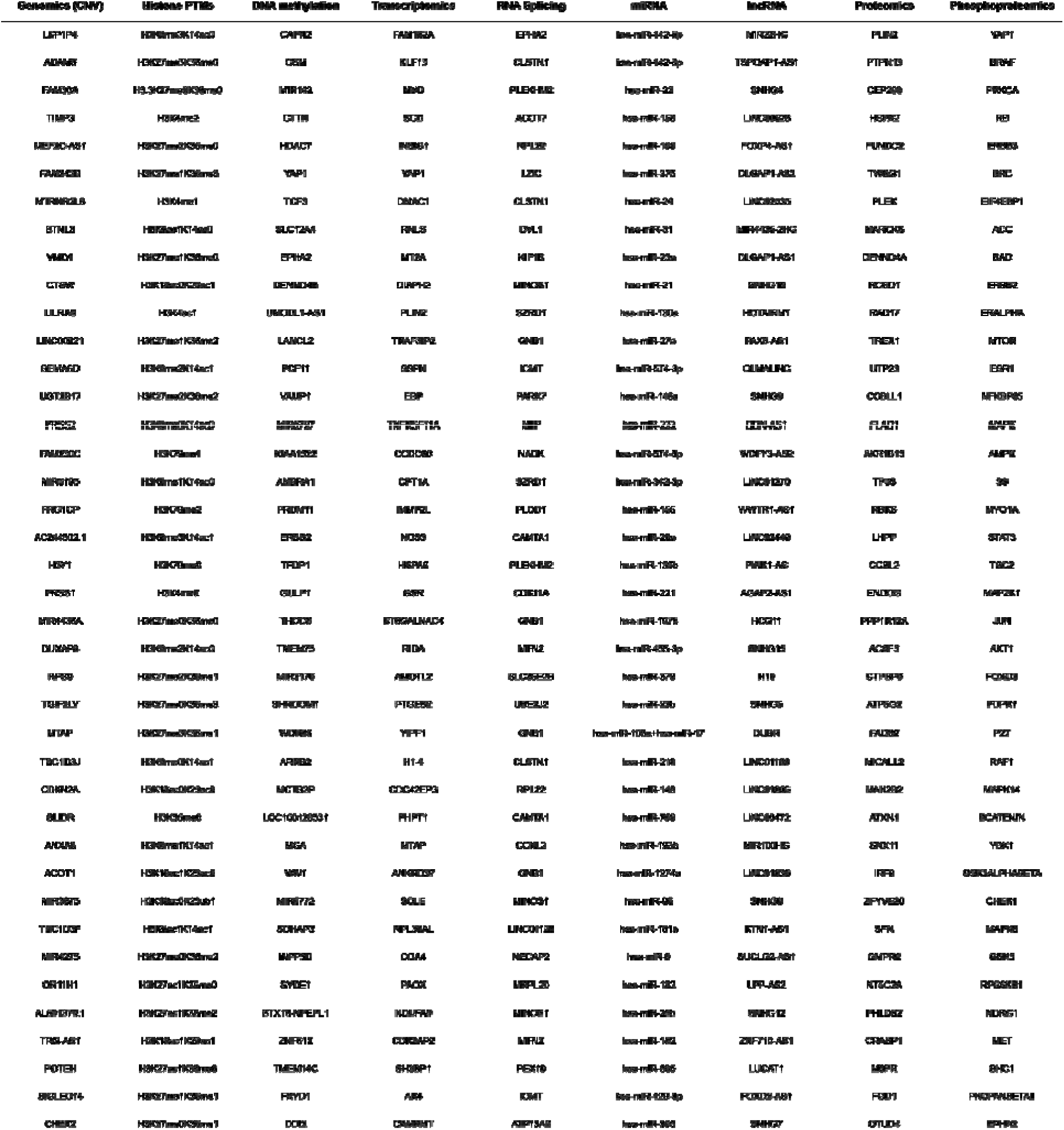
Top 40 global features as identified from 75-25% holdout validation random forests.

**Supplemental Table 3:** Robustness of top features from CCLE transcriptomics in the NCI60 data for full data models and Recon3D genes only. T-test and Kolomogorov-Smirnov test results from the top 1, 3, 5, 10, 50, and 100 CCLE features isolated in NCI60 data as compared to the whole data. Top features for predicting each metabolite were tested against the whole data for the entire metabolite panel (pooled). Data were stacked to create a single vector of feature importance scores across all metabolites for the top feature subsets as well as whole data. Feature importances were used to assess the robustness of the random forest ML framework used to train the CCLE models. Feature importances were significantly higher for all top feature subsets in the pooled analyses using transcriptomics. Top features for individual metabolites also showed significantly higher correlations and feature importance scores for a subset of metabolites (FDR p-value < 0.05). Only using the top 1 feature, CMP, sucrose, GABA, anthranilic acid, N-carbamoyl-beta-alanine, and adenosine displayed conserved top features, which are all in peripheral metabolism. * = significant (p-value < 0.05)

| Top Features | Whole-data |  | Recon3D-only |  |
| --- | --- | --- | --- | --- |
|  | T-test | KS-test | T-test | KS-test |
|  | P value | P value | P value | P value |
| 1 | 2.54E-05* | 6.50E-02 | 3.02E-02* | 5.2E-02 |
| 3 | 2.51E-05* | 7.30E-02 | 2.48E-02* | 6.7E-02 |
| 5 | 2.22E-06* | 4.30E-02* | 1.73E-02* | 2.9E-02* |
| 10 | 4.23E-10* | 2.90E-03* | 4.12E-03* | 2.4E-02* |
| 50 | 1.44E-10* | 4.60E-03* | 8.05E-07* | 4.28E-06* |
| 100 | 2.92E-17* | 3.68E-05* | 3.69E-13* | 1.52E-10* |

**Supplemental Table 4:**
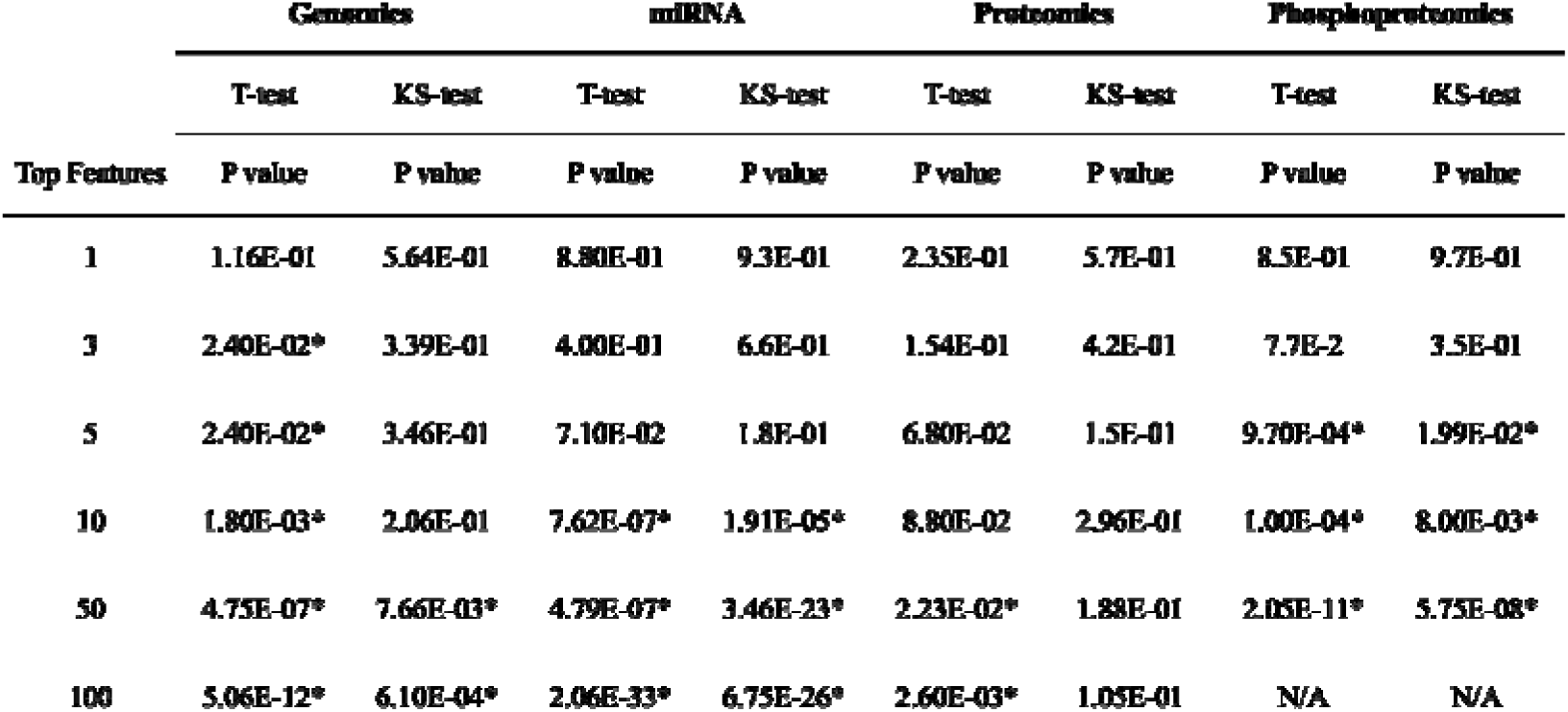
Robustness of top features from CCLE genomics, miRNA, proteomics, and phosphoproteomics in NCI60 models illustrated by T-test and Kolmogorov-Smirnov tests using pooled feature importances. * = significant (p-value < 0.05)

**Supplemental Table 5:** Robustness of top features from CCLE transcriptomics in single-cell models illustrated by T-test and Kolmogorov-Smirnov tests using pooled feature importances. * = significant (p-value < 0.05)

| Top Features | T-test | KS-test |
| --- | --- | --- |
|  | P value (pooled) | P value (pooled) |
| 1 | 2.00E-01 | 4.9E-01 |
| 3 | 2.10E-02* | 1.6E-01 |
| 5 | 6.00E-04* | 2.6E-02* |
| 10 | 6.06E-08* | 3.80E-04* |
| 50 | 8.74E-16* | 3.80E-07* |
| 100 | 2.12E-25* | 4.36E-11* |

**Supplemental Table 6:** Count of occurrences of mutations in the top 20 features (calculated using mutual information) for each of 225 metabolites.

| Mutation | Count |
| --- | --- |
| TP53 (7157) | 20 |
| APC (324) | 19 |
| CDKN2A (1029) | 17 |
| ARID1A (8289) | 17 |
| PTEN (5728) | 16 |
| KMT2D (8085) | 15 |
| TTN (7273) | 14 |
| RB1 (5925) | 14 |
| ACVR2A (92) | 13 |
| MUC16 (94025) | 12 |
| ZNF141 (7700) | 9 |
| KRTAP21-1 (337977) | 8 |
| PCLO (27445) | 8 |
| GPR98 (84059) | 8 |
| RPL22 (6146) | 8 |
| ASXL1 (171023) | 8 |
| NF2 (4771) | 8 |
| CCDC73 (493860) | 8 |
| TEAD2 (8463) | 8 |
| CREBBP (1387) | 8 |

**Supplemental Table 7:** Predicted non-lipid metabolites for which mutations of interest were in the top 20 features.

| Mutation | Predicted Metabolites |
| --- | --- |
| TP53 | CMP, dimethylglycine, kynurenic acid, adenosine, deoxycytidine, carnitine, thymidine, GMP, creatinine, arachidonylcarnitine, adenine, creatine |
| APC | N-carbamoyl beta alanine, betaine, creatine, taurine, NMMA, arachidonylcarnitine, pantothenate, tyrosine, alpha-glycerophosphocholine, phosphocreatine, carnitine, acetylcarnitine, histidine, alpha-hydroxybutyrate, UMP, GABA |
| CDKN2A | Creatinine, NMMA, dimethylglycine, allantoin, UMP, xanthine, hexoses, CMP |
| PTEN | Phenylalanine, carnosine, oleylcarnitine, kynurenic acid, pantothenate, glutathione oxidized, anserine, threonine, leucine |
| RB1 | Isoleucine, lactose, phosphocreatine, phenylalanine, creatine, arginine, adenosine |
| MUC16 | Putrescine, hydroxybutyrate, isoleucine, dimethylglycine, glutathione reduced, creatine, NADP, acetylcarnitine, phosphocreatine |

**Supplemental Table 8:**
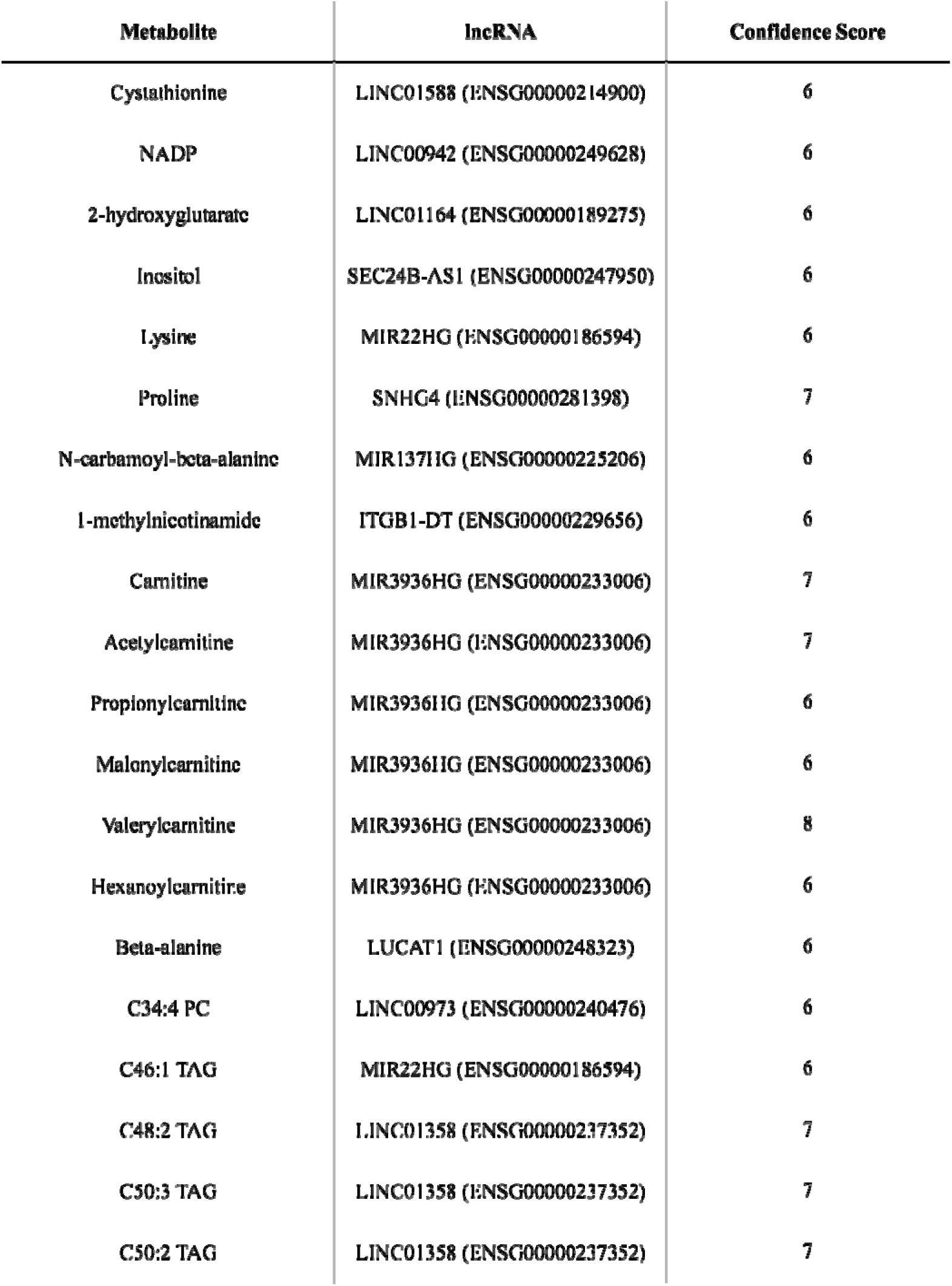
High confidence lncRNAs and their respective predicted metabolite prioritized for further study. lncRNAs were selected based on feature importance (in the top 3 most important features) and confidence score (6 or greater).

| Metabolite | lncRNA | Confidence Score |
| --- | --- | --- |
| Cystathionine | LINC01588 (ENSG00000214900) | 6 |
| NADP | LINC00942 (ENSG00000249628) | 6 |
| 2-hydroxyglutarate | LINC01164 (ENSG00000189275) | 6 |
| Inositol | SEC24B-AS1 (ENSG00000247950) | 6 |
| Lysine | MIR22HG (ENSG00000186594) | 6 |
| Proline | SNHG4 (ENSG00000281398) | 7 |
| N-carbamoyl-beta-alanine | MIR137HG (ENSG00000225206) | 6 |
| 1-methylnicotinamide | ITGB1-DT (ENSG00000229656) | 6 |
| Carnitine | MIR3936HG (ENSG00000233006) | 7 |
| Acetylcarnitine | MIR3936HG (ENSG00000233006) | 7 |
| Propionylcarnitine | MIR3936HG (ENSG00000233006) | 6 |
| Malonylcarnitine | MIR3936HG (ENSG00000233006) | 6 |
| Valerylcarnitine | MIR3936HG (ENSG00000233006) | 8 |
| Hexanoylcarnitine | MIR3936HG (ENSG00000233006) | 6 |
| Beta-alanine | LUCAT1 (ENSG00000248323) | 6 |
| C34:4 PC | LINC00973 (ENSG00000240476) | 6 |
| C46:1 TAG | MIR22HG (ENSG00000186594) | 6 |
| C48:2 TAG | LINC01358 (ENSG00000237352) | 7 |
| C50:3 TAG | LINC01358 (ENSG00000237352) | 7 |
| C50:2 TAG | LINC01358 (ENSG00000237352) | 7 |

**Supplemental Table 9:**
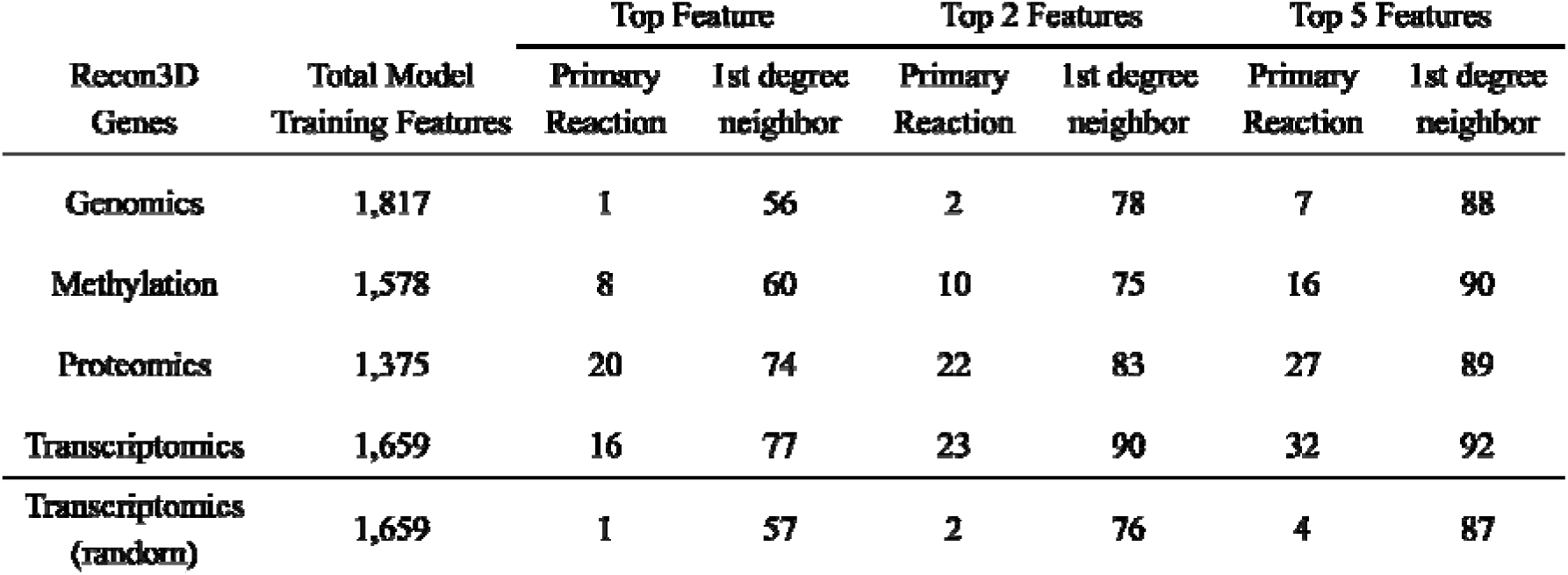
Number of metabolite models trained on the Recon3D metabolic gene subset that had at least one top feature map to Recon3D for the reactions directly associated with or within one neighbor of the predicted metabolite (out of 95 common metabolites). Random analyses (last row) with transcriptomics data involved picking random transcripts per metabolite model and seeing if at least one feature mapped to Recon3D. This process was repeated 50 times, and the median number of overlaps was calculated for each category.

| Recon3D<br>Genes | Total Model<br>Training Features | Top Feature |  | Top 2 Features |  | Top 5 Features |  |
| --- | --- | --- | --- | --- | --- | --- | --- |
|  |  | Primary<br>Reaction | 1st degree<br>neighbor | Primary<br>Reaction | 1st degree<br>neighbor | Primary<br>Reaction | 1st degree<br>neighbor |
| Genomics | 1,817 | 1 | 56 | 2 | 78 | 7 | 88 |
| Methylation | 1,578 | 8 | 60 | 10 | 75 | 16 | 90 |
| Proteomics | 1,375 | 20 | 74 | 22 | 83 | 27 | 89 |
| Transcriptomics | 1,659 | 16 | 77 | 23 | 90 | 32 | 92 |
| Transcriptomics<br>(random) | 1,659 | 1 | 57 | 2 | 76 | 4 | 87 |

**Supplemental Table 10:**
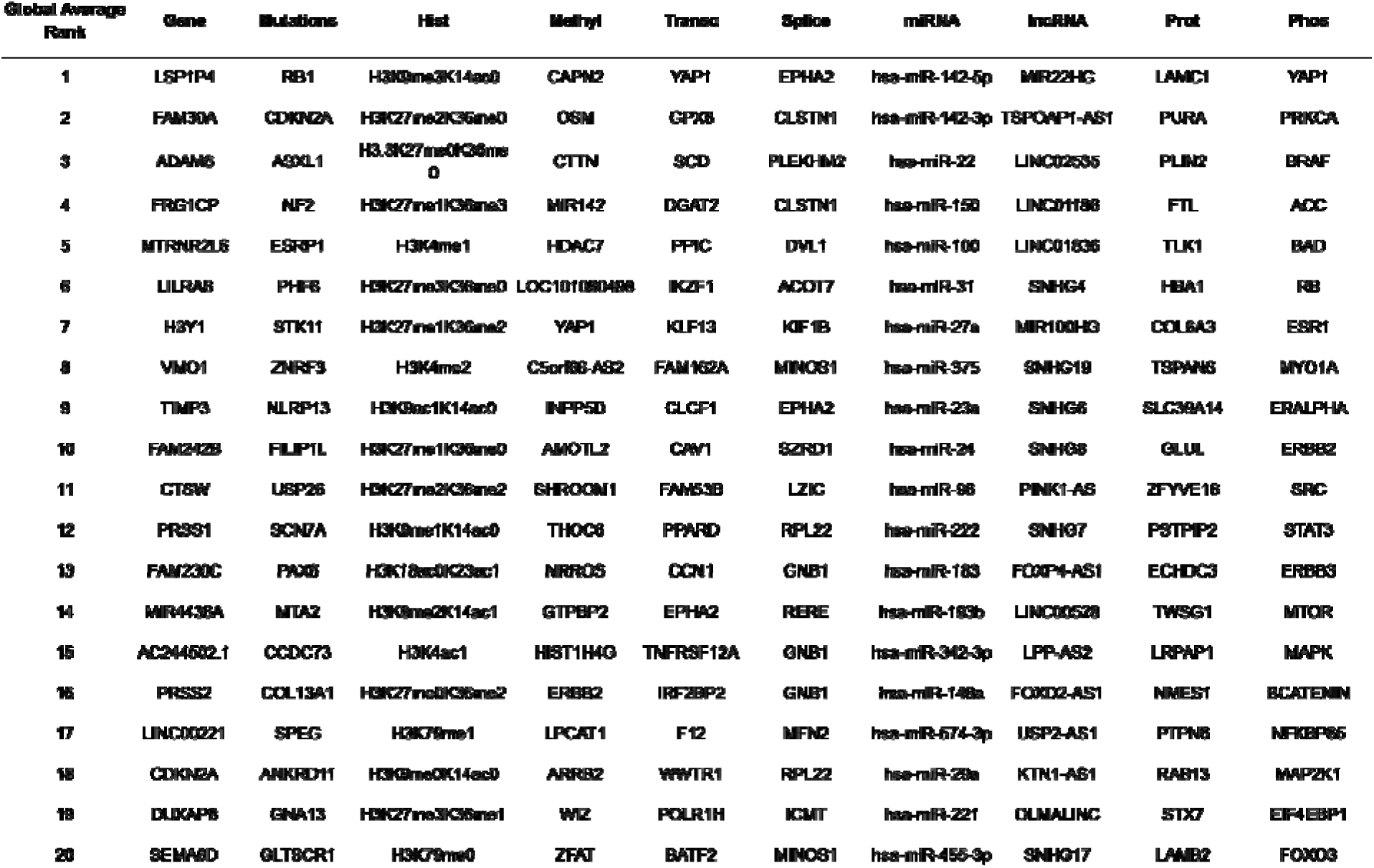
Global feature importance ranks for the top 20 features from each omics class from random forests (except mutations, which used SVMs). Median rank was used to determine the most important features over all 225 metabolite models for each feature.

**Supplemental Table 11:**
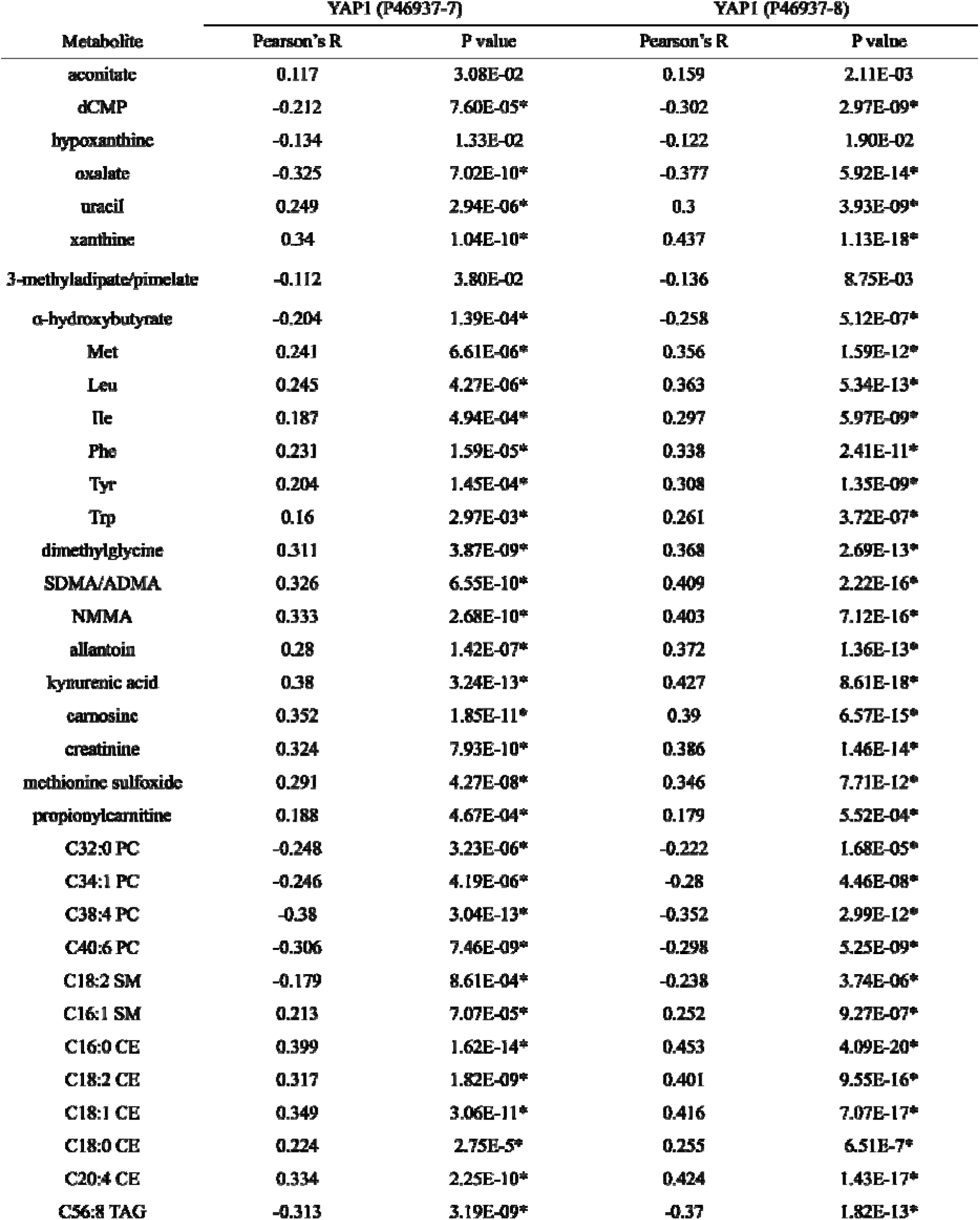
Correlation of two protein isoforms of YAP1 with metabolites for which ML found YAP1 to be the top feature from transcriptomics inputs. Correlations were obtained from proteomic and metabolomic CCLE data using the DepMap portal^46^. Associations between YAP1 and the metabolites it predicted well from ML were assessed for statistical significance using a Bonferroni corrected P value (marked with a *).

## Notes

https://github.com/sriram-lab/Recon8D

https://doi.org/10.7303/syn68236153

### Competing Interest Statement

The authors have declared no competing interest.

### Summary of Updates

We have performed a new analysis across all eight omic regulatory layers in Recon8D and integrated this with known protein-protein interactions, metabolite-reaction, and miRNA-gene interactions from curated databases. This integrated analysis revealed cross-omic interactions and complementary regulatory nodes in the network. We have also incorporated two more omic layers including genomic mutation and lncRNAs, which revealed several well known and novel regulatory interactions. We have further included a novel confidence scoring system based on robustness of interactions across several control experiments. We have also described methods and limitations in greater detail and provided a detailed protocol in our github page.

